# Comprehensive dissection of immune microenvironment in the progression of early gastric cancer at spatial and single-cell resolution

**DOI:** 10.1101/2022.02.16.480776

**Authors:** Tiantian Du, Huiru Gao, Honglei Wu, Juan Li, Peilong Li, Jie Gao, Qiuchen Qi, Xiaoyan Liu, Lutao Du, Yunshan Wang, Chuanxin Wang

## Abstract

While the changes of tumor immune microenvironment (TME) have critical implications for most tumor progression, works that could reveal the compositions and immunity features of TME are needed. Profiling gastric malignant cells at single-cells resolution has shown the transcriptional heterogeneity is represented at different states of gastric cancer, implying that diverse cell states may exist, including immune cells, and all components in TME make some balances in early gastric cancer (EGC) progression. However, it remains unclear how immune cells contributing malignancy of gastritis, constituting general characteristics of gastric TME. Furthermore, the role of specific interactions among cells in gastric TME remains to be illustrated. Here, we performed spatial transcriptomes and single-cell RNA-seq analysis across 18 gastric samples, identifying 17 celltypes and reconstructing their location information. We found that immune cells represented different degree of dysregulations during the progression from non-atrophic gastritis (NAG), atrophic gastritis (AG) to EGC, including imbalance of cytotoxic and inhibitory effects in T cells, maturation inhibition in B cells and malignant genes up-regulated obviously in myeloid cells. Besides, pathway activities showed that hypoxia, reactive oxygen species and fatty metabolism signaling were activated from AG stage, which may accelerate progression of EGC. Moreover, cellular interactions further identified the roles of hypoxia in gastric TME. Overall, the multi-omics data presented in this study offer a comprehensive view of immune cell types, states changes and locations within the gastric tissues during the progression from NAG, AG to EGC, advancing our understanding of the composition and immunity of different gastric states, offering diagnostic and preventive thoughts for EGC.

## Introduction

Gastric cancer possesses the characteristic of high mortality, is the fifth most common cancer. The development of gastric cancer is a multi-step process, generally from non-atrophic gastritis (NAG) to atrophic gastritis (AG) with or without intestinal metaplasia (IM), and finally to all-stage gastric cancer (GC), which is academically known as Correa cascade ^1^. The early gastric cancer (EGC), as initial phase of the process of GC, holds well prognosis more than 90% ^2 3^. Thus, a comprehensive understanding of EGC progression from NAG, on the one hand, is the base of further timely intervention before advanced stages, hopefully improving GC condition ^4^. On the other hand, it is in favor of exploitation of targets for GC risk warning and noninvasive diagnosis, as the most recognized means of EGC diagnosis now remains the invasive endoscopic biopsy ^4 5^.

The progression of most tumors is known as results of dynamic interaction between tumor cells and their immune microenvironment, including EGC ^6^. During the process, gastric mucosa tissues appear heterogeneous changes and malignant cells, which have been reported previously ^7-11^. However, how immune microenvironment paly roles during the EGC progression is still unclear. It was showed that infiltration of inflammatory cells, as well as expression of chemokines and interleukins are different among normal gastric mucosa, simple gastritis, AG and GC ^12^. IL33 was reported to promote metaplasia and its exogenous expression could moderately induce spasmolytic polypeptide-expressing metaplasia (SPEM), may represent as important joint of chronic gastritis and IM ^13^. The typically binary M1/M2 paradigm of macrophages was disappeared, and transcriptionally heterogenous among each cluster was in its place ^14 15^. Besides, 12-gene prognostic signature for metastatic gastric adenocarcinoma (GAC) was established ^16^. Although few works have been reported, they are not enough to illustrate changes during progression from NAG to EGC.

As the complexities of gastric immune microenvironment and technical limitations, previously, it was difficult to profile the component, structure and immunity of immune microenvironment. The recently developed single-cell sequencing (scRNA-seq), VDJ and spatial transcriptomics (ST) sequencing have realized the identity of complicated immune microenvironment. As new techniques, scRNA-seq requires digestive cell suspension and lacks spatially relative location, while ST sequencing represents with spatial spots images with low cell resolution. Then, with the comprehensive use from the dimension of multi-omics, not only the cell biology is elucidated accurately at single cell resolution with scRNA-seq, but the spatial location information and intercellular interaction is acquired by use of ST, even the immunized libraries of T/B cells are obtained ^16 17^. Several tumor and immune microenvironment profiles have been successfully clarified by combined scRNA-seq and ST, including pancreatic cancer ^18^, squamous cell carcinoma ^19^, as well as colorectal cancer liver metastasis ^20^. Thus, we considered application of the combined methods to elucidate microenvironment characteristics during development from NAG, AG to EGC.

In this study, we acquired 21,848 single cells and 11,347 spatial spots. With the combined use of scRNA-seq and ST sequencing, we characterized the dysregulation of gastric immune microenvironment changes. The multi-omics data offer a comprehensive view of immune cell types, states change and locations within the gastric tissues during the progression from NAG, AG to EGC, advancing our understanding of the composition and immunity of different gastric states, offering diagnostic and preventive thoughts for EGC.

## Methods

### Experimental models and subject details

#### Human sample handling (samples and sample information)

Clinical samples were received immediately after extraction from the endoscopy (The delivery time of every sample was less than 30 min) and processed rapidly to ensure viability and quality for scRNA-seq and ST ^21^.

The 12 tissue samples used for ST in our study were randomly selected from established cohort more than 200 specimens, and other 6 fresh tissues used for scRNA-seq were randomly collected from the clinic. All study participants provided written informed consent and study protocols were approved by the ethical review community of the Second Hospital of Shandong University. All of freshly resected biopsy specimens were obtained from one or two sites by conventional upper gastrointestinal endoscopy. Fresh samples were then washed several times with dulbecco’s phosphate-buffered saline (DPBS) for further embedding or digestion ^7^.

#### Pathological annotation for spatial images

All sections used in the spatial study were identified by two independent pathologists. Tissue HE images included tumor cells characteristics were defined as the EGC group, no tumor features but contained intestinal mctaplasia areas were defined as AG group, and tissues with mild inflammatory but no tumor and intestinal metaplasia areas were defined as NAG group.

### Spatial Transcriptomics

#### Slide preparation

For 12 samples used for the ST, trimming them to a suitable size, then embedded with Optimal Cutting Temperature (OCT) embedding solution (Sakura, #4532, USA) and transfer to −80 ℃ refrigerator for quick freezing and long-term storage. Cryosections of the above tissues were cut at 10 μm thickness, 10 consecutive slices for RNA extraction to assess the RNA integrity number (RIN) (RIN ≥ 7 is qualified), and the other 2 consecutive slices for hematoxylin-eosin staining (H&E) staining to confirm the detailed structures of these tissues.

The slides in the Visum spatial gene expression slide and Reagent Kit (10x Genomics, PN-1000184) contained four identical 6.5 × 6.5 mm tissue capture areas, each with ∼5000 spots measuring a diameter of 55 μm and the distance between two spots is 100 μm, which should be expected to encompass 6-10 total cells. An individual spot containing millions of oligonucleotides with features: a partial Illumina TruSeq Read 1 sequence, 16 nt Spatial Barcode (shared within each specific spot), 12 nt unique molecular identifier (UMI) (Identification of duplicate molecules), and 30 nt poly (dT) sequence (captures polyadenylated mRNA for cDNA synthesis).

#### Optimization of the permeabilization time

An optimization slide totally covers eight capture areas, the same size as the GEX slide. six capture areas for cryosections cut at 10 μM thickness, the other two are used for positive and negative controls. An optimal permeabilization time must been identified before a new tissue used for generating Visium Spatial Gene Expression libraries. Briefly, the Visium Spatial Tissue Optimization workflow described as placing tissue sections on six capture areas on a Visium Tissue Optimization slide (from Visium Spatial Gene Expression Reagent Kit, 10x Genomics, PN-1000186). The prepared sections were further fixed, stained, and then permeabilized for different times. The time that resulted in maximum fluorescence signal with the lowest signal diffusion was used as the best permeabilization time, the longer time will be optimal if the signal was same at multiple time points. There were different time points among these tissues, just listed as following: for N22-0, N22-1, N8-1, N21-1, N34-1, C3, C7, C8, C9, 24 min was used; while for N26-0, N28-0, N24-1, 22 min was used.

#### Spatial staining and imaging

Sectioned slides were incubated at 37°C with the active surface facing up for 1 min, and fixed with precooled methyl alcohol in −20℃ for 30 min followed by H&E staining (Eosin, Dako CS701, Hematoxylin Dako S3309, bluing buffer CS702), sections were processed at RT by isopropanol (MilliporeSigma) for 1 min, by hematoxylin (Agilent, Dako S3309) for 7 min, by Bluing Buffer (Agilent, CS702) for 2 min, and by Eosin Mix (MilliporeSigma) for 1 min. Last, slides were incubated for 5 min at 37°C in the Thermocycler Adaptor (10x Genomics, PN-3000380). RNase and DNase free water were used in every washing step. After air-drying, the bright-field images were taken on a Leica DMI8 whole-slide scanner (Germany) at 10× resolution.

#### Permeabilization and reverse transcription

The slides were inserted into slide cassettes (from the from the Visium Slide Kit) to ensure individual reaction chambers of each tissue section. Then according to the predetermined optimal permeabilization time, the tissue sections on the slides were subjected to 70 μL permeabilization enzyme (from the Visium Reagent Kit) at 37℃ for different times to capture polyadenylated mRNA by the primers on the spots. After incubation, each well was washed with 100 μL 0.1×SSC (Sigma-Aldrich), and added with 75 μL reverse transcription Master Mix (from the Visium Reagent Kit) to generate spatially barcoded cDNA.

#### cDNA library preparation and sequencing

After the RT Master Mix was removed from the wells at the end of first-strand synthesis, each well was incubated in 75 μL 0.08 M KOH for 5 min and washed with 100 μL EB buffer (QIAGEN) at room temperature (RT). Then 75 μL Second Strand Mix was added to each well for second-strand synthesis. cDNA amplification and library construction were performed on a S1000TM Touch Thermal Cycler (Bio Rad). Once the Visium Spatial Gene Expression libraries were constructed using Visum spatial Library construction kit (10x Genomics, PN-1000184), further spatial transcriptome sequencing was processed on the Illumina Novaseq6000 platform immediately. The standard paired-end sequencing reached a depth of at least 100,000 reads per spot (pair-end 150 bp, PE150) (performed by CapitalBio Technology, Beijing).

#### Processing of the raw ST data

Raw Visium Spatial RNA-seq data, bright-field and fluorescence microscope images were processed by the Space Ranger software (version 6.1, 10x Genomics) to align reads, filter and construct feature-spot matrixes, UMIs and genes counting, and match spots on the spatial slide images. The GRCh38 v86 genome assembly was used as reference. The QCs were applied before further clustering and gene expression were analyzed by the Seurat R package (version 4.0). Briefly, the spots away from the main section region were discarded. The mitochondrial and ribosomal genes and genes were expressed ≤ 3 spots were filtered. After quantity control, we obtained 11,347 spots for further analysis.

### scRNA-seq

#### Human Gastric tissues dissociation protocol

All gastric tissues were distinguished from non-tumor and tumor regions by gastroscopist before further processing. After several washing with 1×DPBS containing 0.04% bovine serum albumin (BSA, Miltenyi Biotec, 130-091-376), gastric tumor tissues single cells were acquired with digestive enzymes containing 200 μL Enzyme H, 100 μL Enzyme R, 25 μL Enzyme A (all provided in the human tumor dissociation kit (Miltenyi Biotec; 130-095-929), and 4.7 mL DMEM (Gibco; 8117133). While non-tumor tissues digestion was performed in mixed enzymes containing 100 μL Enzyme D, 50μL Enzyme R, 12.5 μL Enzyme A (all provided in the human multi-tissues dissociation kit (Miltenyi Biotec; 130-110-201), and 2.35 mL DMEM (Gibco; 8117133). Single cells were generated from shaking incubation in the water bath at 37 ℃. After the digestive reaction was deactivated by adding another 45 mL fresh DMEM, the cell suspensions were filtered using a 70 μm cells strainer (Miltenyi Biotec; 130-098-462) and then centrifuged at 300g for 5 min. Live nuclear cells were enriched using a Dead Cell Removal kit (Miltenyi Biotec; 130-090-101) and Red Cell Removal kit (Solarbio; R1010). The final cells were re-suspended in 1×DPBS containing 0.04% BSA and prepared for further steps.

#### Library preparation and droplet based scRNA-seq sequencing

Briefly, the droplet based sequencing platform was used in these studies. Gastric tissue suspensions were loaded on a Chromium Single Cell Controller instrument (10x Genomics, Pleasanton, California, USA) to generate single-cell gel beads in emulsion (GEMs). GEM-RT was performed on a S1000TM Touch Thermal Cycler (Bio Rad): 53°C for 45 min, 85°C for 5 min; held at 4°C and stored at −20°C. Then the GEMs were broken and the barcoded, full-length cDNA was amplified, and quality assessed using an Agilent 4200 (CapitalBio Technology, Beijing). scRNA-Seq libraries were prepared using the Chromium Single Cell 5′ Library & Gel Bead Kit (P/N 1000006, 10x Genomics), Chromium Single Cell A Chip Kit (10x Genomics, 120236), Human T Cell (1000005) and Single Cell V(D)J Enrichment Kit, and Human B Cell (1000016) and Single Cell V(D)J Enrichment Kit according to manufacturer’s instructions. The libraries were finally sequenced using an Illumina Novaseq6000 platform with a sequencing depth of at least 100,000 reads per cell with PE150 reading strategy (performed by CapitalBio Technology, Beijing).

### Processing of raw scRNA-seq data

#### Single-cell gene expression quantification, quality control

Alignment, filtering, barcode counting, and unique molecular identifier (UMI) counting of the sequencing data were performed with Cell Ranger software package and mapped to the hg38 reference genome to generate feature-barcode matrix at the base of human reference genome (GRCh38). Then secondary clustering and differential expression analysis of the raw gene expression matrices were further processed with the Seurat R package. Cells with either gene number ranked over the top 1% or fewer than 200, or mitochondrial gene ratio was more than 25% were regarded as unqualified and excluded before downstream analysis. After quantity control, we obtained 28,860 high-quality single cells for further analysis.

#### Integration of multiple samples, batch correction and dimension reduction

We separately used SCTransform (R package, version 0.3.2) and harmony (R package, version 0.1.0) to integrate the ST data and scRNA-seq data from different patients. For scRNA-seq data, the new combined feature-barcode matrix was first applied for normalization, FindVariableFeatures (12000 highly variable genes were identified) and scaling. For spatial data, “SCTransform” function replaced the above procession. Next, principal component analysis (PCA) (RunPCA, from variable genes) was performed. The top 30 principal components (PCs) were selected for FindNeighbors, FindClusters, UMAP (RunUMAP) and TSNE (RunTSNE) to obtain clusters and bidimensional coordinates for each spot or cell and visualization. In this study, the sample source was used defined as batch factor and using “RunHarmony” or “SCTransform” function to reduce the impact of batch effect and generate batch corrected data for downstream analysis.

#### Cell type annotation

Cell type was annotated by referring to published papers manually and singleR (version 1.6.1). For singleR, unbiased cell type recognition from scRNA-seq data was performed by leveraging reference transcriptomic datasets. The known markers used for cell cluster annotation are listed in Supplementary information.

### BCR and TCR analysis

With the 10 x Genomics 5’ V(D)J platform, the immune libraries were built and TCR/BCR sequences were assembled following Cell Ranger VDJ protocol. Assembled contigs were filtered by intersected with barcodes in scRNA-seq. The retained cells were that with at least one TCR α chain (TRA) and one TCR β chain (TRB). For a given T cell, if there were two or more α or β chains assembled, the highest expression level α or β chains was regarded as the dominated α or β chain in the cell. Each unique α-β pairs was defined as a clonotype. T cells with exactly same clonotype were composed a T cell clone.

Similarly, as for BCR clonotype, only cells with at least one heavy chain (IGH) and one light chain (IGK/IGL) were remained. For a given B cell, if there were two or more IGH or IGK/IGL assembled, the highest expression level IGH or IGK/IGL was defined as the dominated IGH or IGK/IGL in the cell. Each unique IGH-IGK/IGL pair was defined as a clonotype. B cells with exactly same clonotype composed a B cell clone.

### Marker gene detection and differential expression analyses

The FindMarkers function with default parameters (logfc.threshold = 0.01) was used to identified different expression genes (DEGs) between two idents. And FindAllMarkers function (wilcox method and logfc.threshold = 0.25 were used) and was called to identified marker genes of each cluster. Genes with adjusted P < 0.05 as the significant ones and reserved.

### Pathway enrichment analysis

The pathway activities of cells were quantified by GSEA, GSVA and AUCell package, respectively. The log-transformed normalization expression matrixes of cells or spots were as inputs and all analysis were performed with default parameters. The referenced gene sets, including 50 hallmark signatures (MSigDB, H sets), 186 KEGG signatures (MSigDB, C2 sets), 1599 Reactome pathways (MSigDB, C2 sets), and 10402 GO signatures (MSigDB, C5, GO sets), were all derived from the Molecular Signatures Database (MSigDB v7.1).

### Trajectory analysis

The pseudotime trajectories were processed both with Monocle (R package, Version 2.20.0) and Monocle 3 (R package, Version 1.0.1) to determine different cell states. The counts data of gene-barcode matrix were used as input. For Monocle, the function of newCellDataSet was used to create a new data object for analysis, then two components and “DDRTree” were used to reduce dimensions, followed by “orderCells”, the results of cell developmental trajectory were presented. The lowerDetectionLimit is 0.5. While for Monocle 3, the new_cell_data_set function was used for objects building, the starting point of the developmental trajectory based on the biological background.

### Cell–cell interaction analysis

cell-cell communication atlases, which contained pairs of ligands and receptors between different cells, were built and visualized by CellChat (R package, Version 1.1.3)^22^. The ligand-receptor interactions included three modules named secreted signaling, ECM receptor signaling, and cell-cell touch signaling, which consisted of multimeric ligand-receptor complexes, soluble agonists and antagonists, as well as stimulatory and inhibitory membrane-bound co-receptors. The min cell number is 3 in filterCommunication function.

### Transcription factor module analysis

R packages including ‘‘SCENIC”, “RcisTarget”, “GENIE3” and “AUCell” were used to analyze the transcription factor of scRNA-seq and ST data. The reference data named “hg38_refseq-r80_10kb_up_and_down_tss.mc9nr.feather” and “hg38_refseq-r80_500bp_up_and_100bp_down_tss.mc9nr.feather” which downloaded from (https://resources.aertslab.org/cistarget/) were used. In the following analysis, we filtered genes, computed a gene-gene correlation matrix, performed transcription factor network analysis and calculated a score for each TF module in each individual cell.

### inferCNV analysis

The CNVs of each cell and each spot were estimated by “inferCNV” package at the basis of gene transcriptome profiles ^23^. For whole cells and spots among three groups (NAG, AG, EGC), the NAG groups were defined as references. Then, for evaluating of somatic hypermutation (SHM), the B cells were defined as controls.

### Spatial Transcriptomics Spots Deconvolution using SPOTlight

SPOTlight was a R package that deconvoluted spots in ST data using scRNA-seq data by Seeded NMF regression ^24^. First, we acquired the marker genes of each cell type in scRNA-seq which was used to deconvoluted the spots in the following by the function called “spotlight_deconvolution”. After adding the percent of cell type in each spot to the metadata of Seurat object, we employed the “SpatialFeaturePlot” to present the cell proportion of ST.

### Correlation analysis

Correlation analysis for different regions, genes or gene sets were all operated by ‘Spearman’ methods of the stat_cor function.

### Multiplex Immunofluorescence

The embedded patient tissues selected randomly from those subjected to spatial transcriptomes sequencing and biobank were sectioned to 6-μm-thick slides. Then the sections were dried for 10 min at room temperature (RT), circled and soaked for 15 min in PBS to removing OCT. After washing 3 times in PBS, the sections were blocked with 10% BSA/PBS for 1 hour at RT with or without washing. The well-blocked sections were then incubated overnight at 4 °C with the first primary antibodies. Next day, after 3 times washing, the sectioned were incubated with the corresponding second antibodies for 2 hours. After 3 times shaker washing, the sectioned were incubated with the second primary antibodies and corresponding second antibodies just repeated steps for the first primary. Lastly, DAPI was used to stained nuclei for 8 min at RT and immunofluorescence images were obtained using digital slice scanners. The primary antibodies used in the assay were listed as following: Anti-Syndecan-1 (1:200, Abcam, ab128936), Anti-pan Cytokeratin (1:200, Abcam, ab7753), and DAPI (Servicebio, G1012).

### Data and code availability

Processed data can be accessed from the NCBI Gene Expression Omnibus (GEO) database when the study acquires permission. The major codes generated during the process will be provided if needed.

## Results

### Exploring the spatial architecture and single cell characteristics of the immune microenvironment in the progression of EGC

To explore the mechanism of transformation from gastritis to EGC, we identified the spatial features of 12 tissue samples from NAG, AG, and EGC patients, separately, which represented different stages in the development of gastric cancer (Figure 1A-C, Supplementary Figure 4A). We recognized distinguishable tumor regions, atrophic regions with or without IM, and regions with no obvious lesions (Figure 1F). Correlation analysis of these regions showed distinctions among the non-lesion regions of each groups much less the distinguishable lesions areas (Supplementary Figure 5A). Then, globally, pathway activities profiles of the spatial regions showed specific activated patterns (Supplementary Figure 1D-E). Comparing with NAG groups, in EGC groups, the MYC signaling, oxidative phosphorylation, interferon alpha response, IL-1 signaling, mTORC1 signaling, TNF signaling, and myeloid leukocyte mediated immunity were up-regulated, while in AG, fatty acid metabolism, xenobiotic metabolism, and lipids metabolism were up-regulated. Inversely, the pathway of KRAS signaling down, aspartic type peptidase activity, B cell receptor signaling, and cell recognition, which up-regulated in NAG groups were all down-regulated in EGC groups (Supplementary Figure 1D-E). Although we found the pathway activities were partly associated with specific celltypes, such as the muscular regions were in line with activated coagulation pathway. For most pathways, their activities were inhomogeneous in each spot and had difficult in matching with particular celltypes, on the one hand for the technical limitations, on the other hand for the complex microenvironment. Besides, there was an evolutionary trend from NAG, AG, to EGC cells analyzed by time trajectory analysis, along with distinct changes of gene expression (Figure 1D-E). Given that, we further explored the difference of cell characteristics and gene expression profiles among three groups with scRNA-seq. We harvested 28,860 cells using 10x scRNA-seq from 6 fresh gastroscopic biopsy specimens of NAG, AG and EGC (Figure 1A). Then we merged, filtered, normalized, batch-corrected, clustered all cells, finally identified 17 cell types of 21,848 cells that passed the quality control (Figure 2A). Epithelium cells, T cells, plasmas occupy a larger portion, fibroblasts and parietal cells have similar number of cells. Besides, there exist endothelium, B cells, mast cells, goblet cells, chief cells and so on, which in a relatively small number (Figure 2E). The distribution of each cell type in different groups and samples was even, indicating no batch effect in the scRNA-seq data (Figure 2B-C). The expression of signature genes shows the correctness of annotation (Figure 2D).

**Figure 1.**
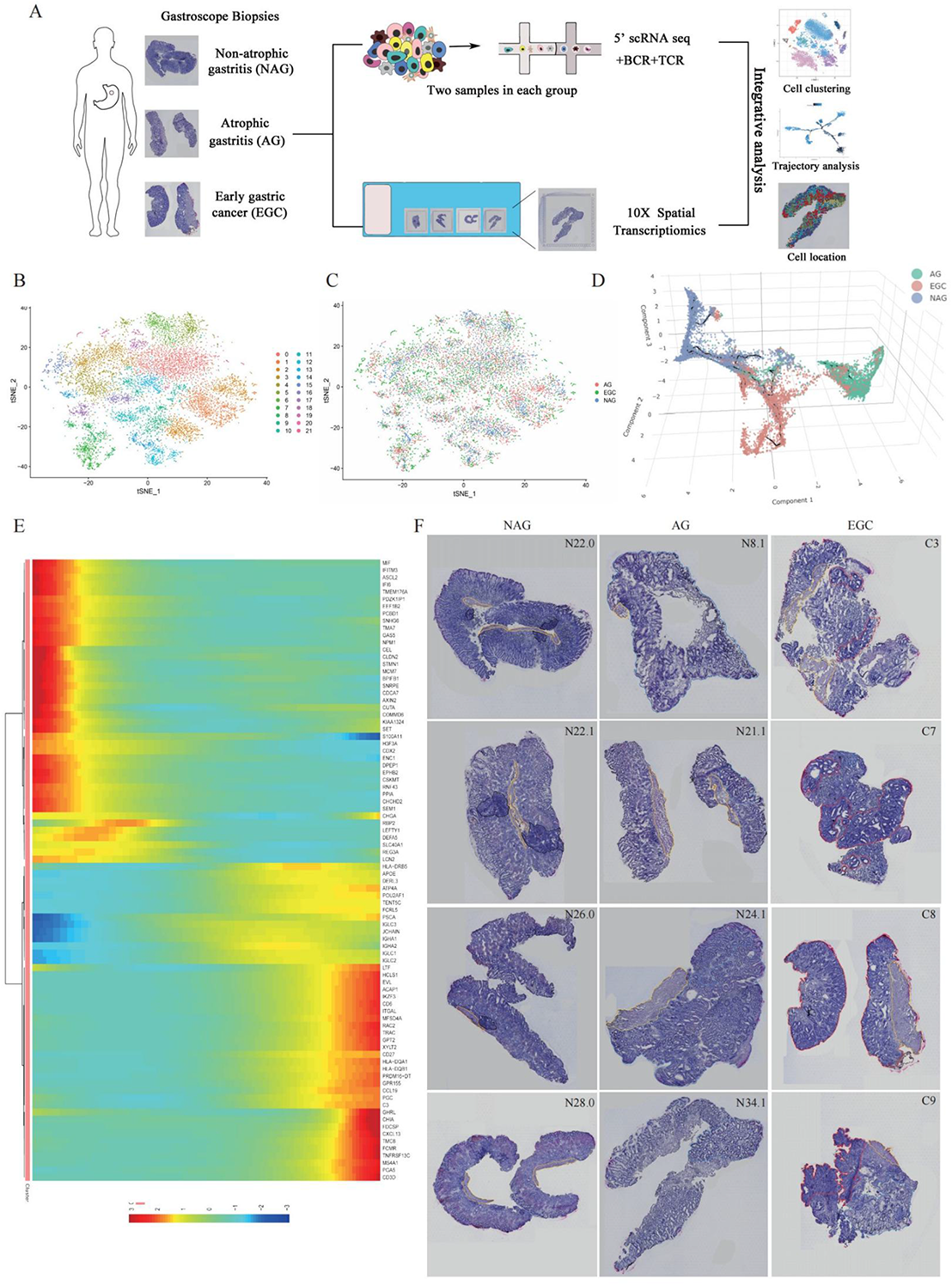
The abstract of spatial transcriptomes sequencing. (A) The flowcharts of scRNA-seq, ST and VDJ sequencing. (B) The TSNE plot for all clusters of ST. (C) The TSNE plot for NAG, AG and EGC three groups of ST. (D) The time trajectory analysis from NAG, AG to EGC of ST. (E) The time trajectory changes of DEGs from NAG to EGC of ST. (F) The images with circled regions of ST sequencing.

**Figure 2.**
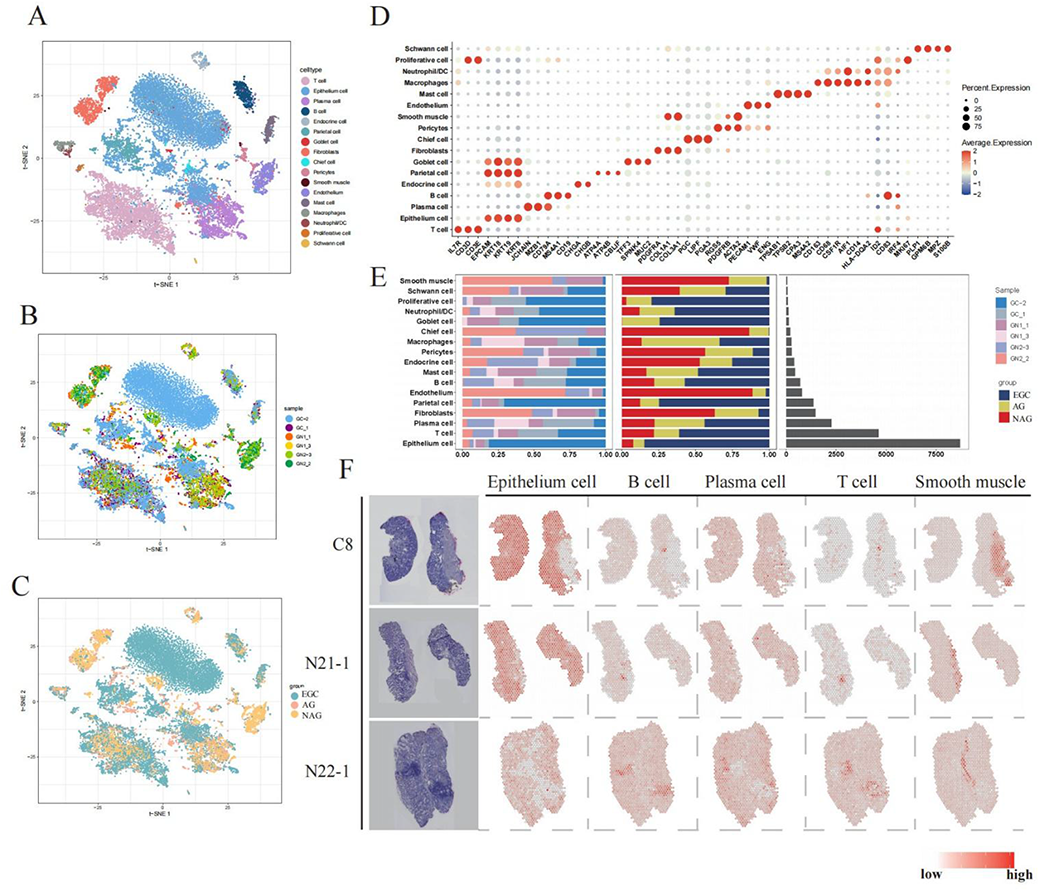
The abstract of single-cell sequencing. (A) The TSNE plot for all identified celltypes of scRNA-seq. (B) The TSNE plot for 6 samples of scRNA-seq. (C) The TSNE plot for NAG, AG and EGC three groups of scRNA-seq. (D) The marker genes for all identified celltypes of scRNA-seq. (E) The cell proportions (left) among 6 samples and 3 groups, as well as cell numbers (right) of all identified celltypes of scRNA-seq. (F) The spatial locations of identified celltypes (epithelium cells, B cells, plasmas, T cells and smooth cells) in 3 ST samples (C8, N21-1 and N22-1).

We explored the origin of each celltype and found considerable heterogeneity between the different groups (Figure 2E). Epithelium cells occupied the largest number in gastric tissues, and most of them were from EGC. Existing in gastric glands, parietal cells were consistent with epithelium cells in specimen sources. For fibroblasts, endothelium cells, and pericytes, there was a decrease from NAG, AG, to EGC, indicating a gradual reduction in vascular and smooth muscle tissues, which may due to sampling differences for biopsy specimens were slightly tiny. For chief cells, the reduction was tally with the signature of SPEM. Furthermore, we explored the expression of TFF2 (Supplementary Figure 6D) which referred in GC progression of SPEM which means SPEM may occurred in our research. In addition, it was worth noting that a group called proliferative cells with the highest proportion in EGC samples, indicating increased ability of proliferation in the development of gastric cancer. Meanwhile, proliferative cells significantly expressed T cell markers such as CD3D and CD3E (Figure 2D), which implied T cell was stimulated in the progression of gastric cancer in proliferation ability.

Then, using SPOTlight, we deconvoluted the ST data combining scRNA-seq data. As a result, we acquired celltype components of each spot. Epithelium cells were widely distributed in spots except for the muscular layer and lymphocyte aggregation area. T cells and B cells were located near each other, and we observed that in the lymphocyte aggregation area, T cells and B cells occupied a relatively high proportion. Compared with the T cells and B cells, plasmas existed in the outer position in the lymphocyte aggregation area. Smooth muscle cells existed mainly in the muscle layer which was reflected in the HE, as it proved the accuracy of cell localization to some extent. The co-localization of Schwann cells, pericytes and fibroblasts indicated that these cells and smooth muscle may be the main cell components of muscle layer. Macrophages tended to exist more at the edge of tissue compared with other immune cells, and the chief cells also avoided the area of lymphocyte aggregation, which was consistent with the epithelium cells. Besides, we also noticed that in NAG, proliferative cells mainly gathered in the lymphocyte aggregation region, but gathered in the epithelial region in the other two groups, suggesting proliferative cells may have different phenotypes and functions including immune cell phenotypes in NAG and Epithelium cell phenotypes in AG and EGC. There also existed interesting phenomenon that the lymphocyte aggregation area was largest in NAG, but diminished in AG and EGC, indicating the immune defense function was the strongest in NAG (Figure 2F and Supplementary Figure 6F).

Previous studied had referred that tumors progression were often accompanied with increased copy number variations (CNV). However, the CNVs had few effects on GC progression, partly given the globally low variations in gastric tissues. In line with the previous work, there were no significant difference of CNV scores during the progression from NAG to EGC (Supplementary Figure 4B-E). The data indicated that there were obvious changes in gene expression profiles and cell characteristics of microenvironment among NAG, AG, and EGC groups.

### Different expression patterns of T cells from NAG to EGC

We noticed that T cells, which play crucial role in tumor microenvironment, accounted for the largest proportion of immune cells. Then, we extracted all the T cells from three groups then reduced the dimensionality and re-clustered them. Depending on gene expression, we divided T cells into eight clusters, including cytotoxic T cells (Cytotoxic T), helper T cells (Th), natural killer cell (NK), double negative T cell (DNT), regulatory T cell (Treg), naive T cell (naive T), exhausted T cell (Exhausted T), as well as a special colony of T cells which expressed high level of epithelial-related genes, called epithium_like T cell (epithium_like) (Figure 3A). T cells from each patient were dispersively distributed (Supplementary Figure 7A). We found large heterogeneity in different samples of T cells. The composition and proportion of T cell showed different characteristics in different disease stages. For example, DNT accounted for larger proportions in NAG and AG stages. It was worth noting that a sample, GN1_1, which belongs to a patient of AG with severe IM, the proportion of T cells was similar to GN1_3, while the latter was similar to AG. To some extent, this proved that AG was intermediate state between NAG and EGC and IM was the state which closed to EGC. For T cell subsets, DNT (CD3+CD4-CD8-T cells) accounted for the highest proportion in the total T cells. Cytotoxic T and exhausted T cells only existed in the samples of EGC and AG, and the proportion of DNT cells gradually decreaseed during the progression of the disease. Naïve T, Th, NK cells showed an increasing trend (Figure 3B, Supplementary Figure 7B). Meanwhile, we used classical markers to characterize the correctness of the clustering. As can be seen, the marker genes presented specific expression in specific T cells (Figure 3C).

**Figure 3.**
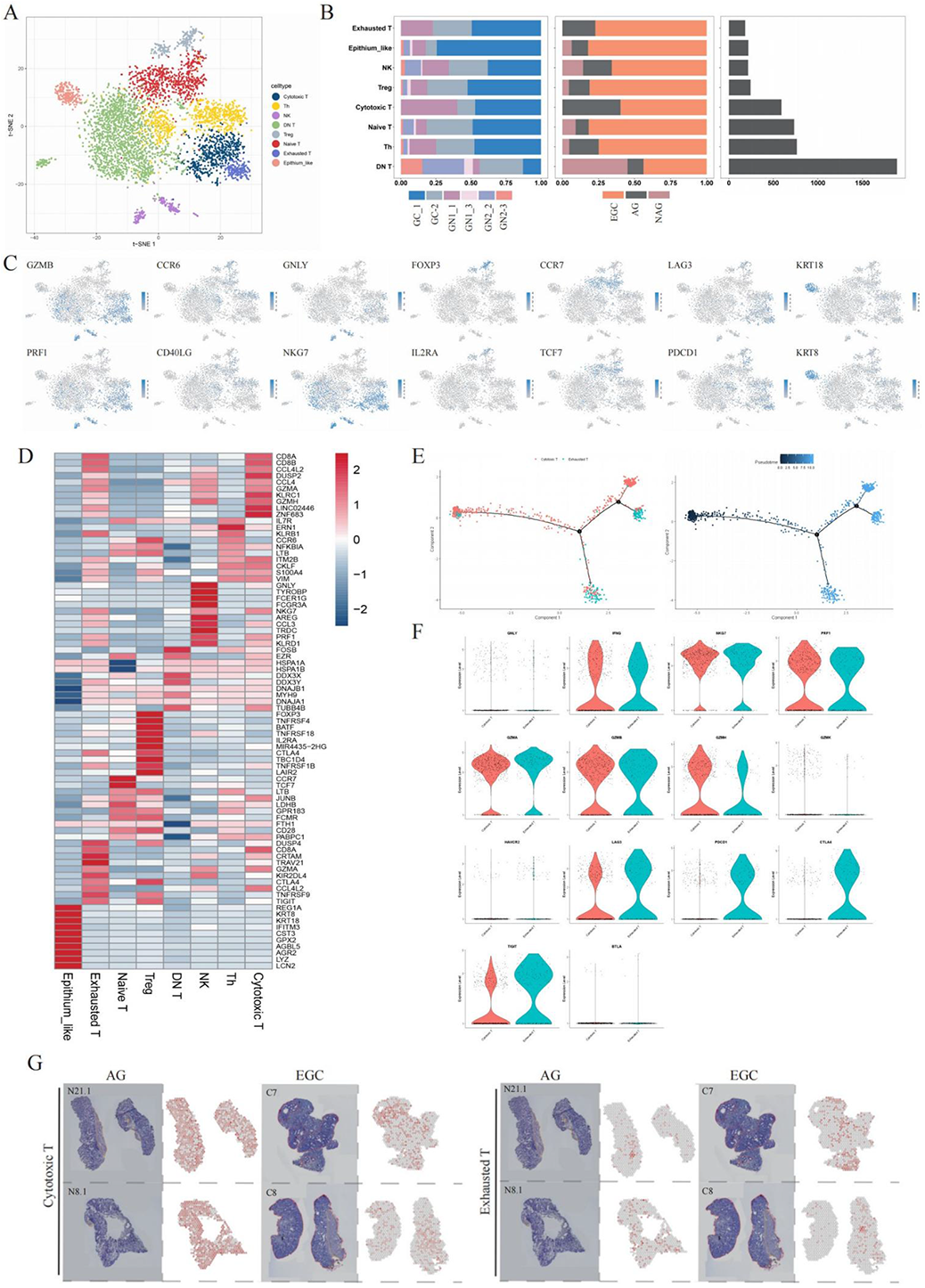
The characteristics of subclusters in T cells. (A) The TSNE plot for subclusters of T cells. (B) The cell proportions (left) among 6 samples and 3 groups, as well as cell numbers (right) of subclusters of T cells. (C) The TSNE feature plots for marker genes of subclusters in T cells. (D) The top 10 marker genes of each subcluster of T cells. (E) The time trajectory between Cytotoxic and exhausted T cells. (F) The relative expression of markers genes in Cytotoxic and exhausted T cells. (G) The spatial locations of Cytotoxic and exhausted T cells in AG (N21.1 and N8.1) and EGC (C7 and C8) groups.

We showed the top 10 DEGs of T cell subsets using heatmap (Figure 3D). The expression of DEGs in heatmap further verified the accuracy of annotation. For cytotoxic T cells, in addition to cytotoxic genes, there were also high expression of KLRC1 ^25^, ZNF683^26^ that interacted with NK cells. There were certain similarities in gene expression of cytotoxic T cells and exhausted T cells. But DUSP4 were higher in exhausted T cells while DUSP2 in cytotoxic T cells. KIR2DL4 which activated cytotoxicity of NK cells were high in exhausted T cells, indicated that both cytotoxic T cells and exhausted T cells could interact with NK cells. DNT concentrated on heat shock protein-related genes, including HSPA1A ^27^, HSPA1B^28^, DNAJB1^29^, DNAJA1^30^. DDX3X is related to the formation of NLRP3 inflammasome ^31^. EZR ^32^ and MYH9 ^33^ were in connection with cell polarity such as migration and invasion.

We tracked analysis of total T cells and found no obviously developmental pattern. However, when we just focused on cytotoxic T cells and exhausted T cells, we found the cytotoxic T cells were prior to exhausted T cells and most of the exhausted T cells gathered in the terminal of the branch as a whole, implying the conversion of these two kinds of cells (Figure 3E). We compared the expression of functional genes between cytotoxic T and exhausted T cells. For cytotoxic genes including IFNG, NKG7, PRF1, GZMA, GZMB, GZMH, there was no difference between the two celltypes. But for immune checkpoint including LAG3, PDCD1, CTLA4 as well as TIGIT, expression of the exhausted T cells was remarkably higher (Figure 3F). The same cell cluster may exhibit diverse states in distinct patients, by finding DEGs, we got the number of cytotoxic T and exhausted T cells DEGs in AG and EGC group.

The total pathway activation patterns were similar between cytotoxic T cells and exhausted T cells, while the latter powered higher activities, except intestinal immune network for IgA production, WNT beta-catenin signal, DNA methylation signaling were mainly enriched in exhausted T cells. Comparing cytotoxic T cells and exhausted T cells in AG groups, the cells in EGC groups presented higher activity in pathways including TRAF6 mediated IRF7 activation pathway, RUNX3 pathway, INF signaling, CXC chemokine receptor signaling and immune response to tumor cells (Supplementary Figure 8B).

Then we used SPOTlight to locate cytotoxic T cells and exhausted T cells. The distribution of the two types of cells was similar to some extent, but cytotoxic T cells had a wider location (Figure 3G, Supplementary Figure 7F).

When analyzing subsets of T cells, we found that a group of cells that neither CD4 nor CD8 was expressed which we called DNT (Supplementary Figure 8C-E). Meanwhile, the number of DNT was 1890, accounting for 39% of T cells and 8.7% of all cells. Although there were some reports about DNT, the function and effect of DNT were still not clear, and hitherto there was no cover about DNT in GC, then we focused on the DNT in the following.

We explored the difference between the three groups in DNT. The trajectory suggested DNT changed step by step as the disease develops (Supplementary Figure 8F). We checked the gene expression of DNT over pseudotime, and the heat shock protein-related genes such as HSPA9, HSPA1A, HSPA1B, HSP90AB1 as well as DEAD-box helicase genes such as DDX5, DDX3X, DDX3Y were taper off (Supplementary Figure 9A). On the contrary, the phenotype of CD8+ T cells gradually emerged, as the CD8A, GZMK were upregulated. In the later periods of pseudotime, the genes of JCHAIN and IG-related genes highly expressed in plasma cells (Supplementary Figure 9B). The gene expression of DNT pseudotime indicated that transitions to CD8+ T cells and plasmas may occur in later development of DNT.

SPOTlight showed that DNT distributed pervasively (Supplementary Figure 8D, Supplementary Figure 9C). But interestingly, areas of lymphocyte aggregation gathered fewer DNT compared to other regions. In the sites of cancer lesions, there existed relatively more DNT (Figure 3G, Supplementary Figure 7F). In combination with the results of monocle, we wildly guessed that DNT may be converted into CD8+ T cells to exert the function of inhibiting cancer at the lesion site (Supplementary Figure 8F).

### The dysregulation of B cells development during EGC progression

In contrast to T cells, B cells population showed less number and diversity between subgroups (Figure 4A-B). According to the unique marker genes, these cells were divided into B cells (CD19, CD20/MS4A1, FCER2 and MHC Ⅱ molecules) and plasmas (TNFRSF17, SLAMF7, XBP1, SDC1 and IGHs) (Figure 4E-F). In line with the previous studies that gastric tissues contained massive plasmas, we also found the phenomenon which hitherto with unknown mechanisms^34^. Spatial location also indicated the diffuse distribution of plasmas while scarce B cells (Figure 4G, Supplementary Figure 11A-D). However, the cell numbers of B cells and plasmas is comparable, respectively, among NAG, AG and EGC groups (Figure 4C-D).

**Figure 4.**
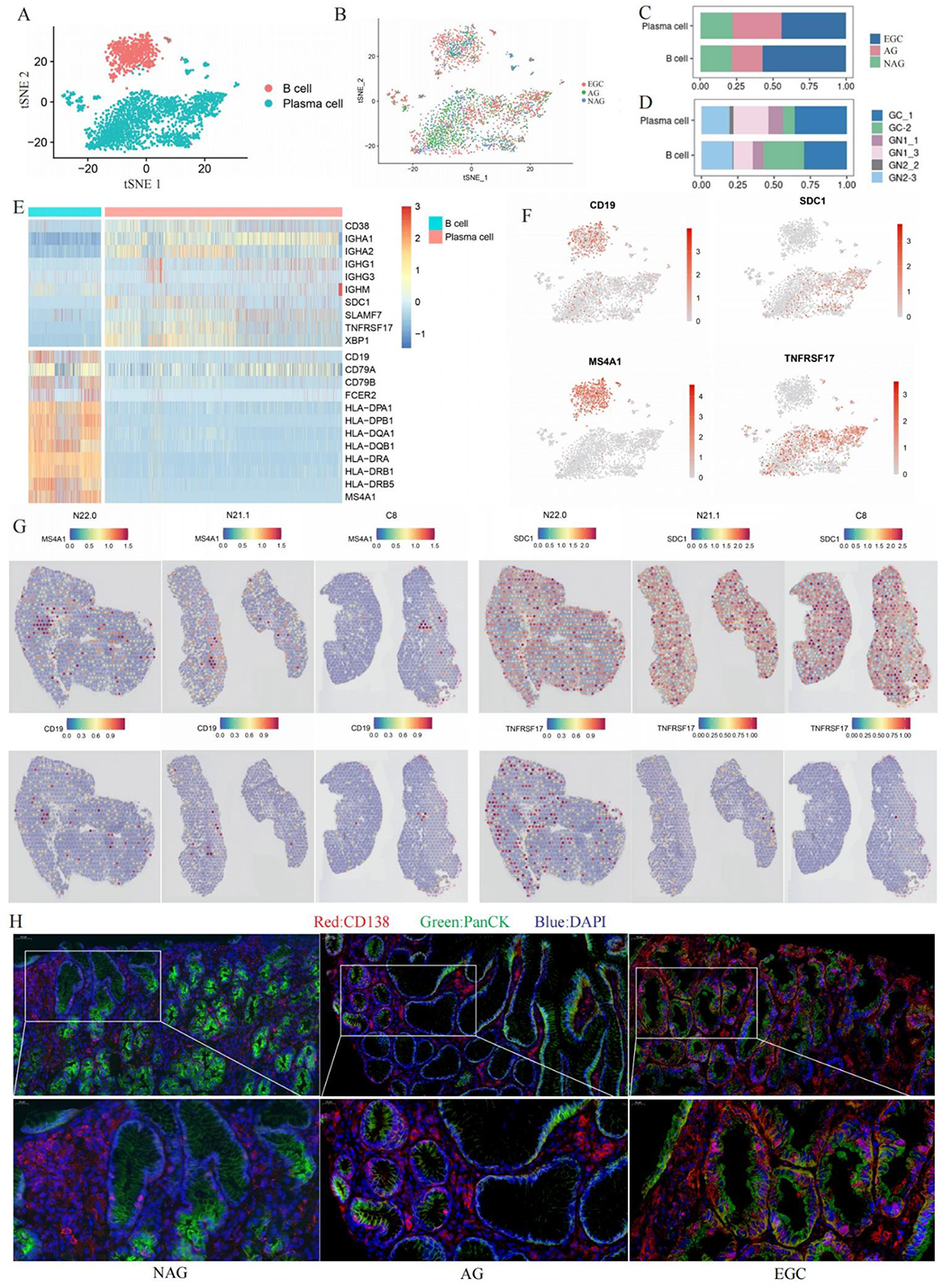
The dysregulation of B cell subclusters. (A) The TSNE plot for B cells and plasmas. (B) The TSNE plot for B cells and plasmas among 3 groups. (C) The cell proportions among 3 goups of B cells and plasmas. (D) The cell proportions among 6 samples of B cells and plasmas. (E) The marker genes of B cells and plasmas. (F) The TSNE feature plots of markers genes of B cells and plasmas. (G) The spatial location of marker genes of B cells and plasmas. (H) The IF staining of plasmas and epithelium cells (Red: CD138, represented plasmas; Green: PanCK represented epithelial cells; Bule: DAPI, represented cell nucleus).

Higher resolution clustering showed there were 7 different subgroups in B cells (Supplementary Figure 10A). B-c0 and B-c5 mainly originated from non-EGC groups while in B-c1, B-c2, B-c3, B-c4 and B-c6, cells from EGC and AG groups predominated (Supplementary Figure 10C). Gene expression analysis showed the similar up-regulated genes between B-c0 and B-c5, DDX5 ^35 36^, DDX3X^37^ and NCL ^38^ were all anti-pathogens and anti-tumor genes. As plenty of oncogenes significantly up-regulated in B-c1 cluster, we inferred it was deteriorating cell populations induced by tumor cells. Then, pathway activity analysis indicating the highly up-regulated activity of pro-tumor signaling, which also showing the malignant potentials of these cells (Supplementary Figure 10F). For B-c2 cluster, RGS13 ^39^, MEF2B^40^, GCSAM ^41^ and LRMP ^42^, essential genes for development of germinal center, were up-regulated. However, FCRL4, usually expressed on the memory B cells under infection, it’s overexpression also could impede antigen-induced BCR signaling ^43^. Besides, ACTG1 ^44^ and FAM3C ^45 46^, which were reported to promote tumor development, were also up-regulated in this cluster. For B-c3, it was defined as follicular B cells for the expression of IGHD and FCER2. TCL1A ^47^, TNFAIP3 ^48^, BTG1 and TXNIP ^49^, pro-tumor or pro-inflammatory genes, were also expressed in this cluster. For B-c4, it was defined as memory B cells, and the marker gene FABP5 was reported to regulate fatty acid metabolism ^50 51^. In B-c6, SIK2 was reported to regulate metabolism reprogramming through PI3K/AKT/HIF axis, promoting tumor progression; WDR76 could activate fatty metabolism restriction ^52 53^; CNR2 may inhibit antibody response and is reported as central effector of B cells restriction of glucose and energy supply ^54^, which in turn indicting that this cluster was lined with the metabolism changes of B cells (Supplementary Figure 10F).

TF analysis identified IRF8, PAX5 and SPIB, the well-known TF essential for development of B cells, CREB3L2 and XBP1 for maturation of plasmas. Moreover, transcriptional activity of EGR3, KLF12, MECP2, HDAC1 and SOX5 were higher in c0-mermory B cells and c5-naïve B cells. On the contrary, FOXP1, HMGB1, EGR1 and BCL11A were lower in these clusters but higher in other B cell clusters. It was reported that FOXP1 presented physiological reduction in germinal center B cells, implying that the high activity of FOXP1 leading to infantilization of B cells ^55^. HMGB1 could be activated after stimulation of hypoxia or inflammation ^56^. BCL11A could exert functions in early stages of B cells ^57 58^. EGR3 could regulate proliferation of B cells through antigen receptor signaling and controlling inflammation ^59^. HDAC1 could regulate the G1-to-S phase transition of cell cycle, its ablation blocked B cells at early stages ^60^. SOX5 was important for development of memory B cells ^61^ (Supplementary Figure 10E).

With further dissection of plasmas, we found these cells possessed the evolutionary trajectory from plasmablasts, shorter-term plasmas to longer-term plasmas (Supplementary Figure 10B, 10D, 10K). Given somatic hypermutation (SHM) and class switch during affinity maturation, plasmas could obtain more variations than B cells, and inferCNV analysis also indicted these differences (Supplementary Figure 10G-J).

Interestingly, spatial feature images showed the diffuse distribution of plasmas and epithelial cells in gastric tissues, on the other hand, the clear clustering was difficult between the two celltypes. Thus, IF was used to figure out the spatial location of them. We found considerable numbers of neighborhood relationships between the two celltypes, especially in EGC patients (Figure 4H). However, there were no illustration for the above phenomenon^34^, which need further exploration.

As TCR clonotypes were different between NAG, AG and EGC patients, we also identified the BCR information (Supplementary Figure 12A). Diversity of clonotypes were increased from NAG, AG, to EGC groups, (Supplementary Figure 12B-C). Besides, the V-J pairs whatever Heavy chains or Light chains among three groups were unique and different (Supplementary Figure 12D-E).

### Characteristics of myeloid cells in the TME of EGC

We then dissected the gene signatures of myeloid subsets revealed in this study (Figure 5A, Supplementary Figure 13A-B). Two mast cell clusters expressed a canonical gene pattern, such as TSPAB1, TPSB2, CPA3, and KIT (Figure 5D, 5G, Supplementary Figure 14C-D), and further distinguished by unique expression of PGA3, NEAT1, HSP90AB1 and TNFAIP3, AREG, MIF, respectively (Figure 5D). Notably, previous studies showed upregulation of TNFAIP3 could promotes gastric cancer development ^62^, and AREG could enhance suppressive function of Tregs ^63^. The two clusters showed different enrichment in EGC, AG, and NAG groups (Figure 5B). Mast c0 cluster existed in all three groups more or less evenly, while mast c1 nearly disappeared in NAG groups. The specific distribution may imply different biological roles they functioned. Pathway scores showed mast c0 cluster correlated with KRAS signaling down, intestinal absorption, ABC transporters, and Hedgehog signaling pathway. Mast c1 cluster tended to become a malignant cell population, and significantly enriched with PI3K-AKT-mTOR, ɑ-TNF, fatty acid metabolism, hypoxia, inflammatory response and IFN-γ signaling (Supplementary Figure 13C).

**Figure 5.**
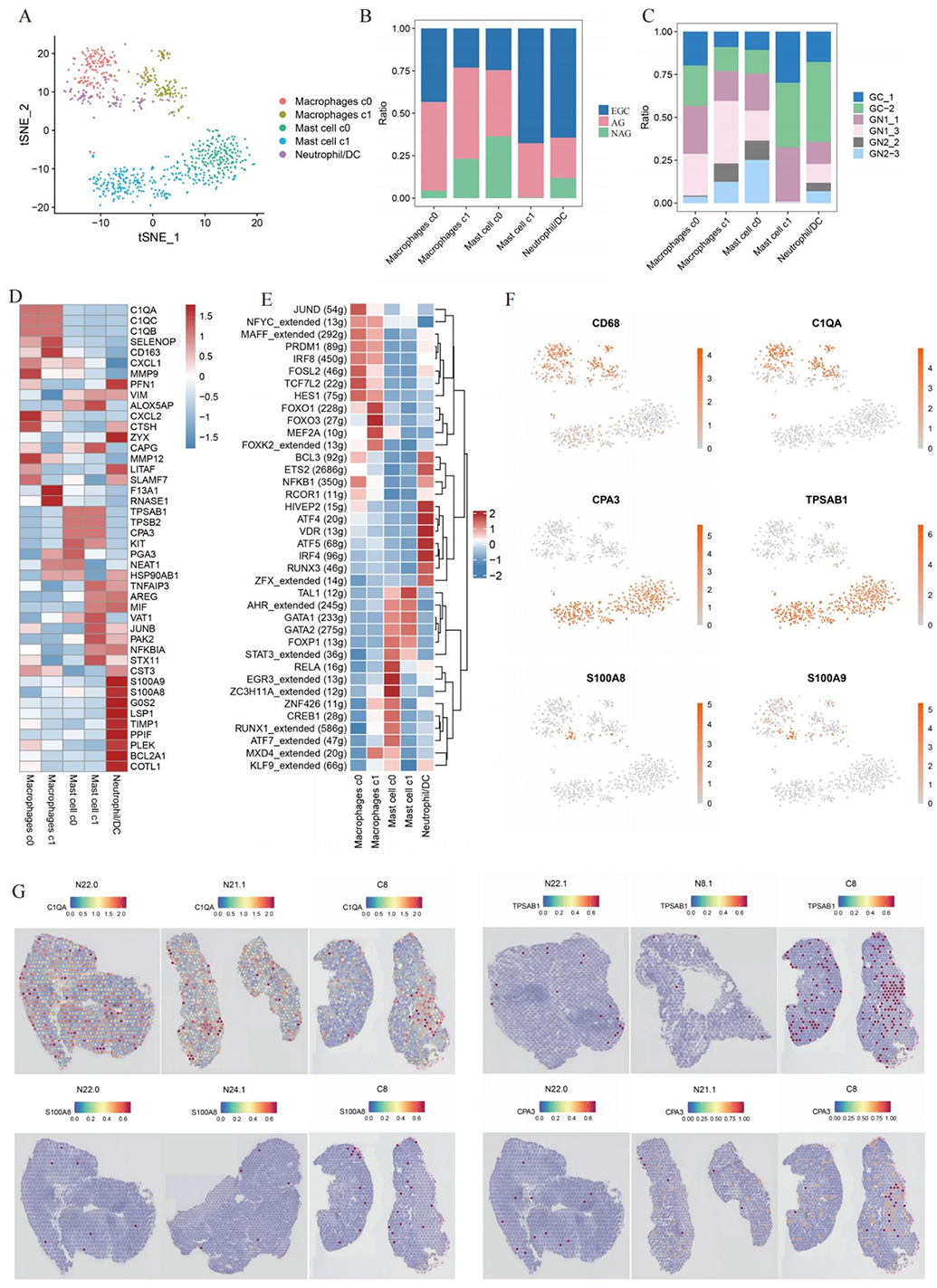
The features of myeloid cells. (A) The TSNE plot for myeloid cells. (B) The cell proportions among 3 goups of myeloid cells. (C) The cell proportions among 6 samples of myeloid cells. (D) The marker genes of myeloid cells. (E) The transcriptional factors of myeloid cells. (F) The TSNE feature plots of markers genes of myeloid cells. (G) The spatial location of marker genes of myeloid cells.

TF analysis indicated that JUND, MAFF, PRDM1, FOSL2, TCF7L2 and HES1 were activated in macrophages c0 cluster, while the TF activity of FOXO1, FOXO3 and MEF2A were higher in macrophages c1 cluster. As for mast cells, AHR, GATA1, GATA2 were well-known TF, while RELA, EGR3 and ZC3H11A were higher in mast c0 cluster. HIVEP2, ATF4, VDR, ATF5 and IRF4 hold higher activity in Neutrophil/DC cells (Figure 5E).

Two macrophage clusters were identified based on their specific expression of C1QA, C1QB, C1QC, and CD163 (Figure 5D, 5G, Supplementary Figure 14A). Among them, clusters showing marked (macrophage c0 cluster) or slightly (macrophage c1 cluster) skewed distribution in three groups (Figure 5B-C). Expression of MMP9, CXCL2, MMP12 and CTSH were observed in macrophage c0 population, both of which exhibited lower expression in macrophage c1 population (Figure 5D). The remaining clusters were Neutrophil/DC population, characterized by high expression of S100A8, A100A9, G0S2 and LSP1 (Figure 5G, Supplementary Figure 14B). Most of the cluster originated from EGC patients (Figure 5B), and the gene expression as well as activities of enriched pathways were also partial to pro-tumor (Supplementary Figure 13C). Both of macrophages and DC population demonstrated the characteristics of antigen presenting cells (APCs), with abilities of antigen procession and presentation. Apart from this, cell adhesion molecules cams, intestinal immune network for IgA production and complement coagulation cascades signaling were also activated in these cells (Supplementary Figure 13C).

### The ligand-receptor interactions of immune cells along with EGC progression

As for the complicated and high-frequency interaction in tissues microenvironment, especially in tumor tissues. We further evaluated the cell interaction among NAG, AG, and EGC groups. The interaction strength and numbers for secreted signaling, cell to cell contact signaling and ECM receptor signaling were all different among three groups. As contrast to NAG group, the interaction numbers and strengths between AG and EGC groups were comparative, indicating AG patients contained conditions driven early tumor occurrence (Figure 6A, 6C-D, Supplementary Figure 15E-F, G, Supplementary Figure 16A, Supplementary Figure 16A-B). Difference analysis of ligand-receptors indicated that LGALS9_CD44, LGALS9_CD45, LGALS9 -HAVCR2(TIM-3), MIF-CD74/CD44, MIF-CD74/CXCR4 and MDK-NCL were major interactions (Figure 6B, Supplementary Figure 15A-D, Supplementary Figure 16C-E, Supplementary Figure 16C-D). We found LGALS9 was up-regulated in mast cells, macrophages, parietal cells and goblet cells. LGALS9_CD45 reported to inhibit B cell receptor signaling ^64^. LGALS9_CD44 could enhance stability and function of Tregs ^65^. Besides, LGALS9 could interact with HAVCR2 (TIM-3) to protect breast cancer against cytotoxic killing of T cells ^66^, partly by inducing apoptosis or exhaustion of effector T cells ^67^. TIM-3, was reported as a specific surface molecule of Th1 cells and could mediate Th1 immune responses ^68^, its dysregulation could lead to inhibited inflammatory response even tumor evasion ^69^. It could promote CD8+ T cell exhaustion and induce production of myeloid-derived suppressor cells (MDSCs)^70^. In our study, TIM-3 mainly expressed in myeloid cells, in line with the previous reports that TIM-3 expression was predominantly localized to myeloid cells in both human and murine tumors ^71^. TIM-3 in mast cells may induce apoptosis ^72^. Macrophage migration inhibitory factor (MIF), an upstream regulator of adaptive and innate immune, but if dysregulated, was a key driver of acute or chronic inflammation and functioned in leukocyte recruitment ^73^. MIF was occurred in varieties of immune cells among AG and EGC groups. CXCR4, one of receptor of MIF, is highly expressed in EGC group, especially in B cells and T cells population. It was reported that CXCR4 could impeded T-lymphocyte infiltration and immunotherapy in other tumors ^74^. CD74/CD44, as the receptor complex of MIF, its activation could trigger multiple intracellular signal pathway, including ERK pathway, PI3K/AKT signal, NFκB and MAPK pathway ^75^. As the hypoxia pathway was activated in gastritis conditions, it was reported that HIF-1α-and HIF-2α-dependent MIF could regulate function of myeloid cells partly by interacting with CD74/CXCR4 complexes, and then activate the p38/MAPK and PI3K-AKT signals ^76^. On the other hand, the complex could also function as positive regulators of B cells homeostasis and maturation ^77^. Midkine (MDK) was reported to support progression of gastric cancer ^78^, partly by inhibiting expression of Granzyme B (GZMB) and cytotoxicity of NK cells ^79^, as well as increased epidermal growth factor receptor (EGFR) signaling under hypoxia through interaction with surface nucleolin (NCL) ^80^. It was also identified to contribute to multidrug resistance in GC cells ^81^. Besides, MDK rewires immunosuppressive environment in melanoma and gallbladder cancer ^82 83^. In this study, we found that MDK is overexpressed in EGC and AG patients comparing with NAG ones. One of receptors of MDK was NCL, a major nucleolar protein of growing cells, also functioned as a cell surface receptor to shuttle between cytoplasm and nucleus, thus providing a mechanism for extracellular regulation of nuclear events ^84^.

**Figure 6.**
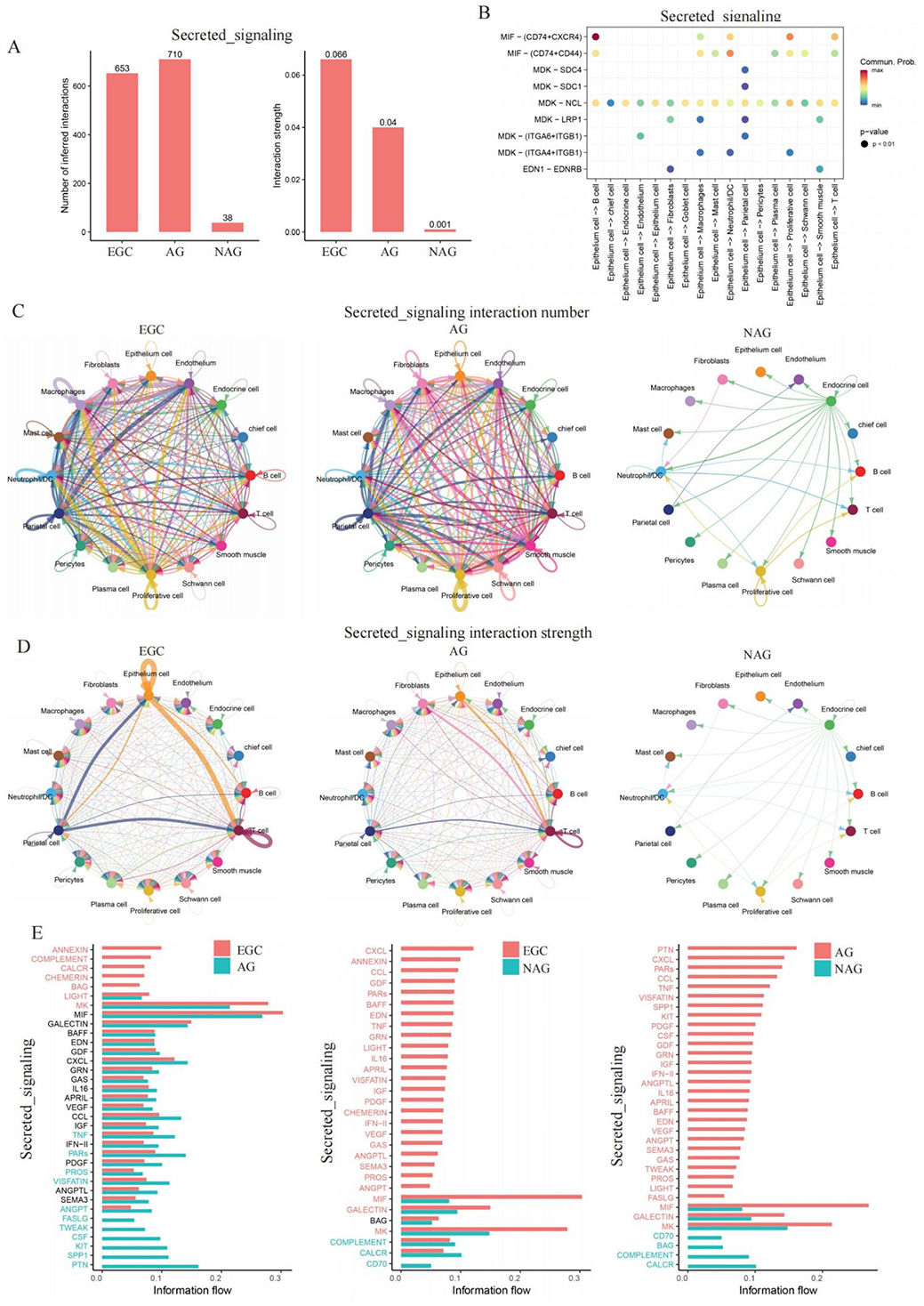
The cellular interaction among identified celltypes. (A) The cellular interaction numbers and strengths among 3 groups. (B) The ligand-receptor pairs from epithelium cell to other celltypes. (C) The cellular interaction numbers of secreted signaling. (D) The cellular interaction strengths of secreted signaling. (E) The interaction involved pathways between AG and EGC (left), NAG and EGC (median), as well as NAG and AG (right) of secreted signaling.

## Discussion

In order to explore the immune microenvironment changes from gastritis to early gastric cancer, we firstly dissected the immune cells component and spatial structures by comprehensively use of single-cell sequencing and spatial transcriptome technologies. Although there were serval gastric cancer-related studies previously, they either focused on the malignant cells, or other un-immune cells in gastric cancer just at single cell resolution ^7-11^ ^85^, or with other spatial methods such as DSP instead of ST^34^.

Global pathway enrichment and spatial regions analysis indicated that several pathways such as hypoxia conditions, reactive oxygen species pathway and fatty metabolism were activated in AG stages, all benefited to the progression of EGC and form of inhibitory immune microenvironment, comparing with the NAG group. While MYC, INFα, IL-1 and TNF signaling, all tumor-related pathways were activated in EGC groups. The overall changes implied the heterogeneity of the compositions. Then in this study, multiple immune cells, including T cells, B cells and myeloid cells were found and they possessed different states and expression patterns among EGC, AG and NAG patients.

When the gastric disease progressed from NAG to EGC, the balance between effector and inhibitory states of T cells were impaired gradually. Specifically, exhausted T cells and cytotoxic T cells were emerged in AG and EGC stages, which were non-existent in NAG state, and immune responses activated obviously. This was consistent with the previous work that early tumor progression was accompanied with immune response activation. In addition to the existence of exhausted T cells, another natural inhibitory role in TME, Tregs, up-regulated transcriptional activity of HIVEP1 and STAT3 in NAG stage, while enhanced activities of IRF4 and IKZF2 in AG state, and possessed higher transcriptional activities of FOXP3, BATF, and PRDM1 in EGC period. Among them, HIVEP1 was a newly found TF in Tregs, it was just reported highly activated in TNFRSF9+ Treg cells, at a late part of the trajectory ^86^, presenting a different pattern from our results. As previous reports, PRDM1 usually up-regulated transcriptional activity in advanced or end-stage, implying worse immune effects ^87 88^. These changes of transcriptional factors activity during progression from NAG to EGC may indicate that the inhibitory effects of T cells were one of the contributing factors of early gastric cancer. The above also showed that the exhausted or inhibitory immune responses have begun from AG patients, implied the preventive measures for EGC maybe should enforce from this stage. Besides, we found a rare celltype in T cells, the DNT cells, which occupied a large portion but has uncertain marker genes to defined them, which was in accordance with the previous reports ^89-91^. Time trajectory showed the continuous evolution of DNT from NAG, AG, to EGC, without enough experimental verification, thus, the significances of their existence were unclear, which was worth further investigation in the future.

Then, the same as T cells, dysregulation also emerged in B cells. The higher transcriptional activity of FOXP1 and BCL11A in clusters (B-c1, B-c2, B-c3, B-c4 and B-c6) that enriched more in AG and EGC groups, represented the infantilization of B cells ^55 57 58^. Besides, the transcriptional inhibition of SOX5 indicated the loss of mature memory B cells ^61^, as well as maturation disorders for the lower activities of EGR3 ^59^ and HDAC1^60^, which positively regulate cell cycles. Comparing B cells, plasmas were rich in gastric tissues, the same as previous reports ^34^. We have found an interesting phenomenon that plasmas were co-localized or accompanied with epithelial cells, especially in EGC groups, which confirmed by ST images of marker genes expression and IF staining. However, the cellular interaction could not find out hints for the phenomenon, showing possibilities for further exploration in depth.

There were two malignant clusters, Mast c1 and macrophage c0, mainly enriched in AG and EGC groups, which expressed high level genes that promoted tumor progression and inhibited effective immune responses. As for AREG up-regulated in Mast c1, could interact with EGFR to enforce the function of Tregs ^63 92^. LGALS9, mainly expressed in myeloid cells, could interact with TIM-3 to exert immune suppressive roles ^66 69^. Besides, both MIF-CD74/CXCR4 and MDK-NCL could activate in hypoxia condition, and regulate functions of T cells or myeloid cells, respectively ^76 80^.

The strength of this study came from application of combined technologies: spatial transcriptomics, scRNA-seq and VDJ analysis. Visium ST has allowed us to determine the location of cell types within different tissues; although scRNA-seq could not acquire enough depths of gene expression just like the common RNA-Seq, it has allowed us to confirm the known celltypes in immune microenvironment during progression of gastric cancer, as well as separate rare cell types such as Schwann cells and DNT cells.

However, there were also several limitations in this study: (1) Given the samples incorporated in this study were not enough, more samples were needed. (2) We mainly focused on the immune cells, other un-immune cells which were also extremely important for EGC progression, reported previously ^7-11^, were not clarified in detail, it will be presented in latter works. (3) The presented conclusions mostly depended on the computationally analysis, and more functional experiment were needed to convey for further identification. (4) The exploration of molecular mechanisms was not enough, which needed more experimental outcomes as bridges to every hypothesis.

Overall, we characterized the dysregulation of immune microenvironment changes both in single-cells and spatial transcriptome resolution, the multi-omics data presented in this study offer a comprehensive view of immune cell types, states change and locations within the gastric tissues during the progression from NAG, AG to EGC, advancing our understanding of the composition and immunity of different gastric states, offering diagnostic and preventive thoughts for EGC.

## Acknowledgement

We thank Department of pathology in the Second Hospital of Shandong University for species and technical assistance.

---

**Supplementary Figure 1.**
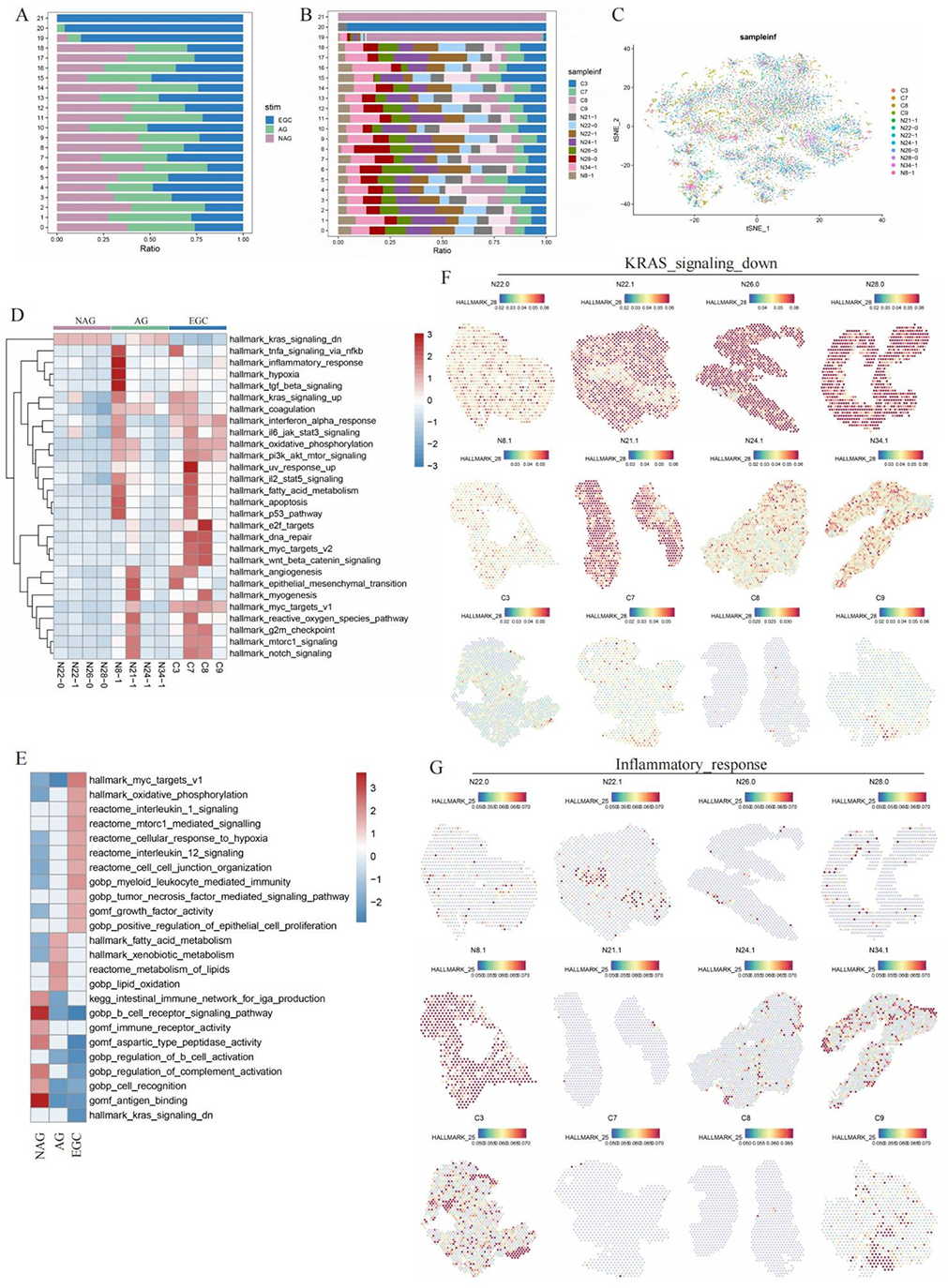
The pathway activities of samples used for ST, related to Figure 1. (A) The cell proportions among 3 groups of ST data. (B) The cell proportions among 12 samples of ST data. (C) The TSNE plot for ST data among 12 samples. (D) The key pathways enriched in each sample used for ST. (E) The key pathways enriched in each group of ST. The pathway activities of (F) KRAS down signaling down pathway and (G) Inflammatory response pathway represented by spots among 12 sample used for ST.

**Supplementary Figure 2.**
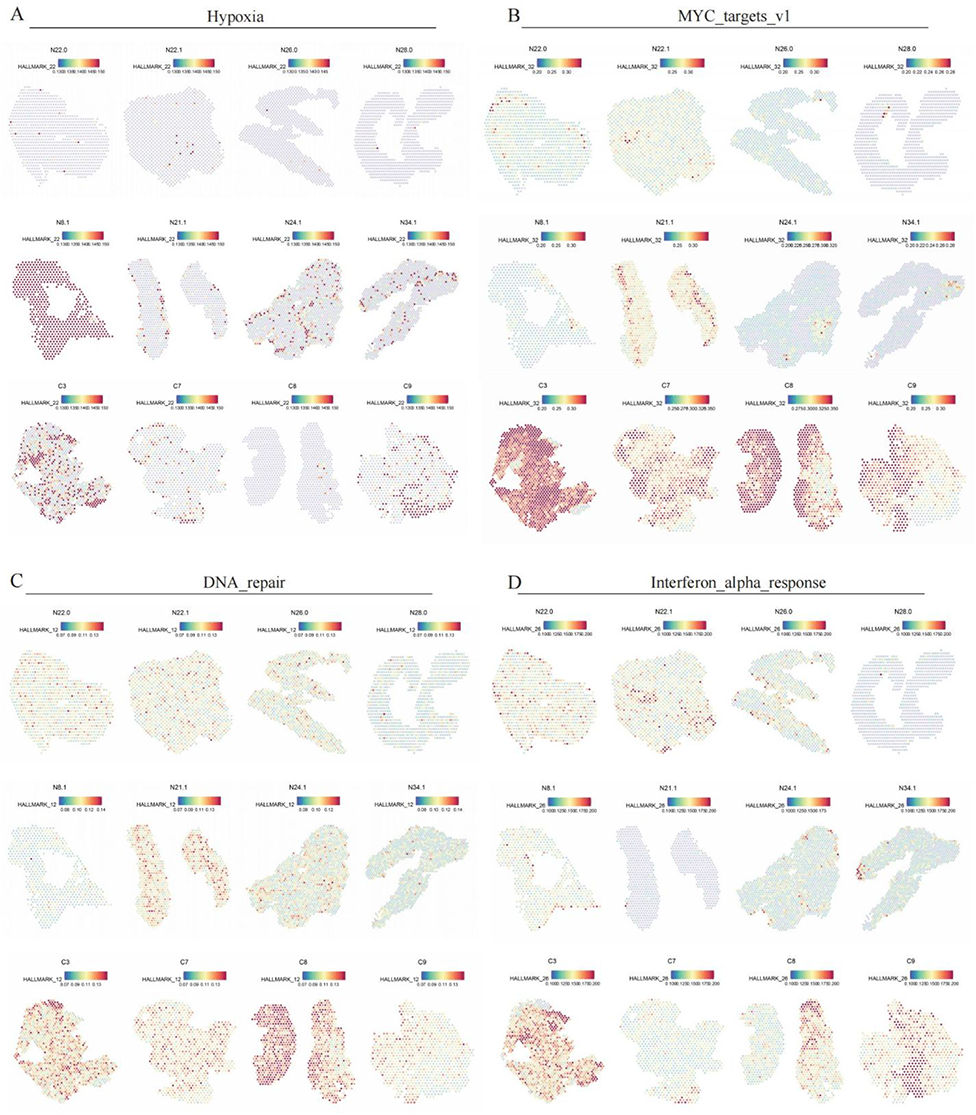
The activities changes of 4 key pathway in 12 samples used for ST, related to Figure 1. The pathway activities of (A) Hypoxia, (B) MYC signaling, (C) DNA repair, and (D) Interferon alpha response pathway represented by spots among 12 sample used for ST.

**Supplementary Figure 3.**
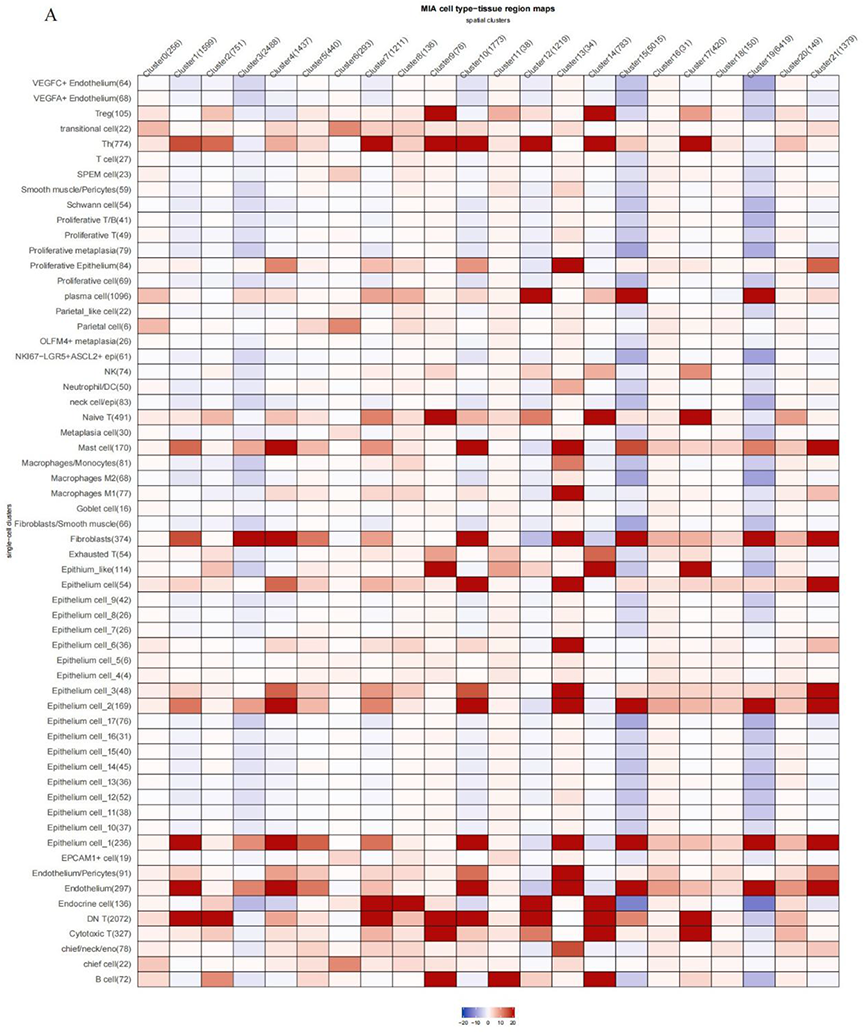
The identification of celltypes by MIA methods, related to Figure 1. (A) The identification of celltypes by MIA methods.

**Supplementary Figure 4.**
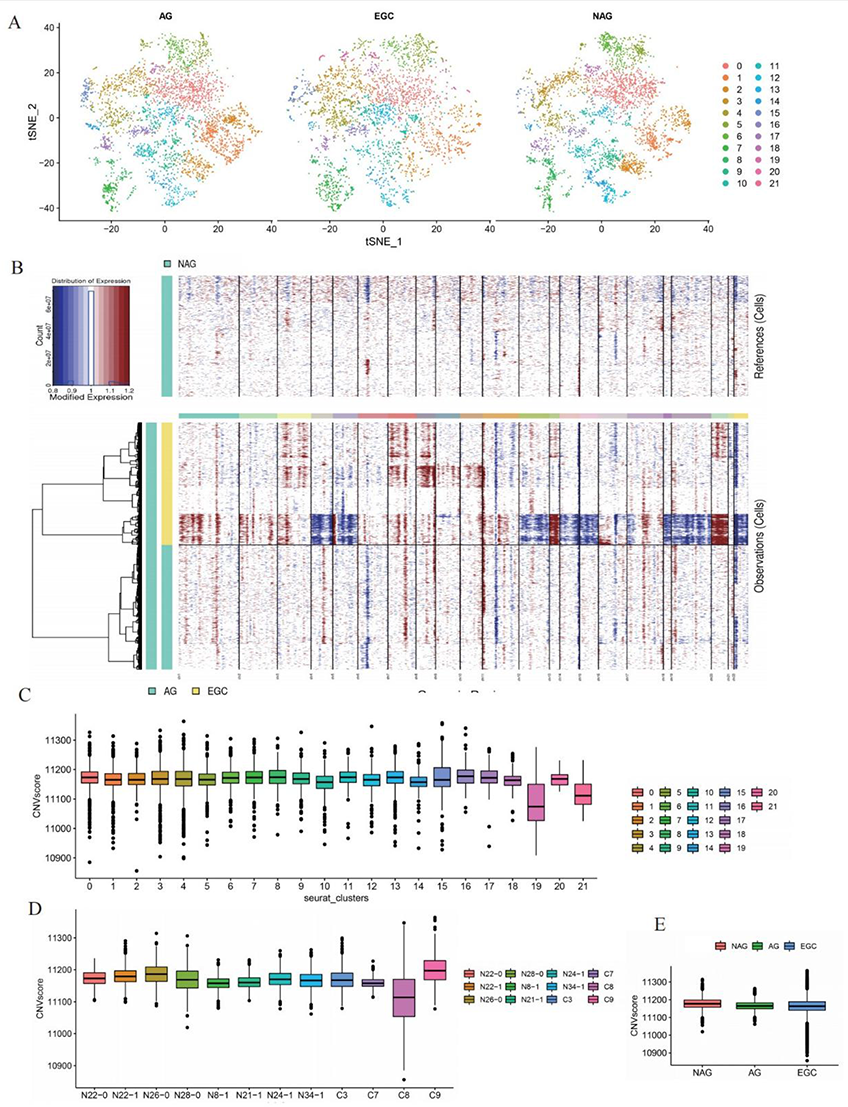
The evaluation of CNV in gastric TME, related to Figure 1. (A) The TSNE plot of all clusters of ST data split by 3 groups. (B) The evaluation of CNV changes among 3 groups by inferCNV. The NAG group was used as control. (C) The CNV score in each cluster of ST data. (D) The CNV score in each sample of ST data. (E) The CNV score in each group of ST data.

**Supplementary Figure 5.**
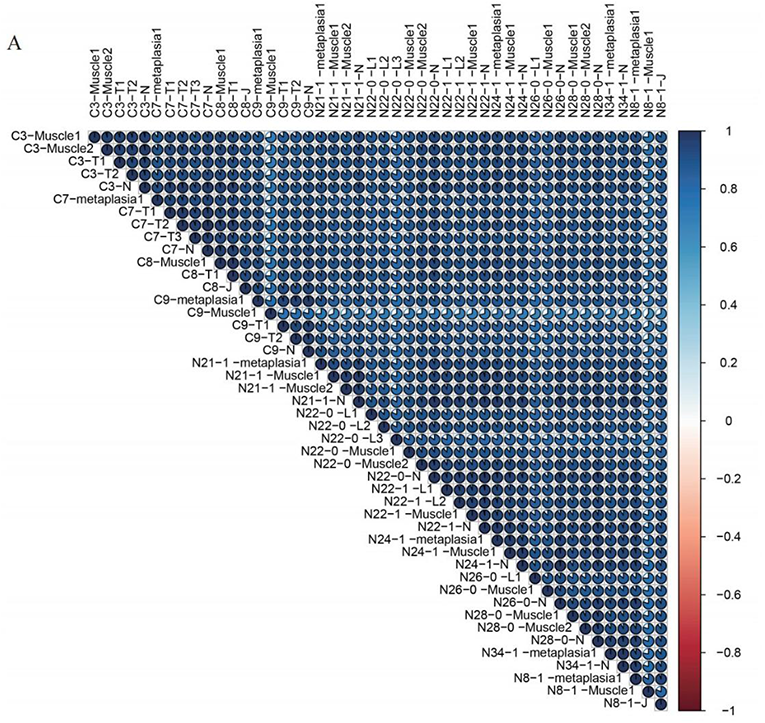
The similarity of specific regions in each ST image, related to Figure 1. (A) The similarity of specific regions in each ST image.

**Supplementary Figure 6.**
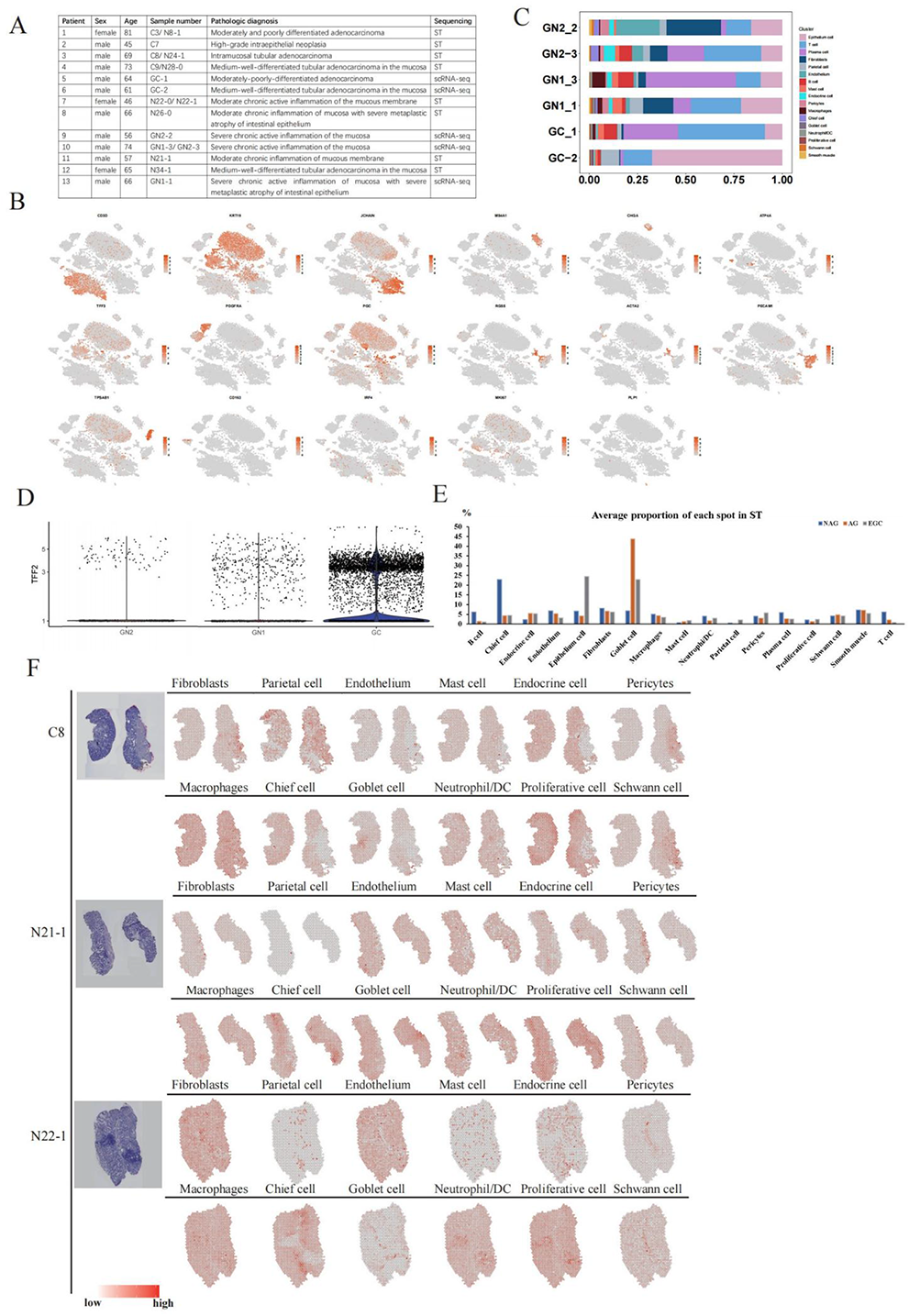
The abstract of single-cell sequencing, related to Figure 2. (A) The pathological information of patients used in this study. (B) The TSNE feature plots of marker genes for identified celltypes. (C) The cell proportions of identified celltypes among 6 patients used for scRNA-seq. (D) The expression of TFF2 among NAG, AG and EGC groups. (E) Average proportion of each spot in identified celltypes. (F) The celltype location in spatial images (C8, N21-1 and N22-1) of fibroblasts, parietal cells, endothelium, mast cells, endocrine cells and pericytes.

**Supplementary Figure 7.**
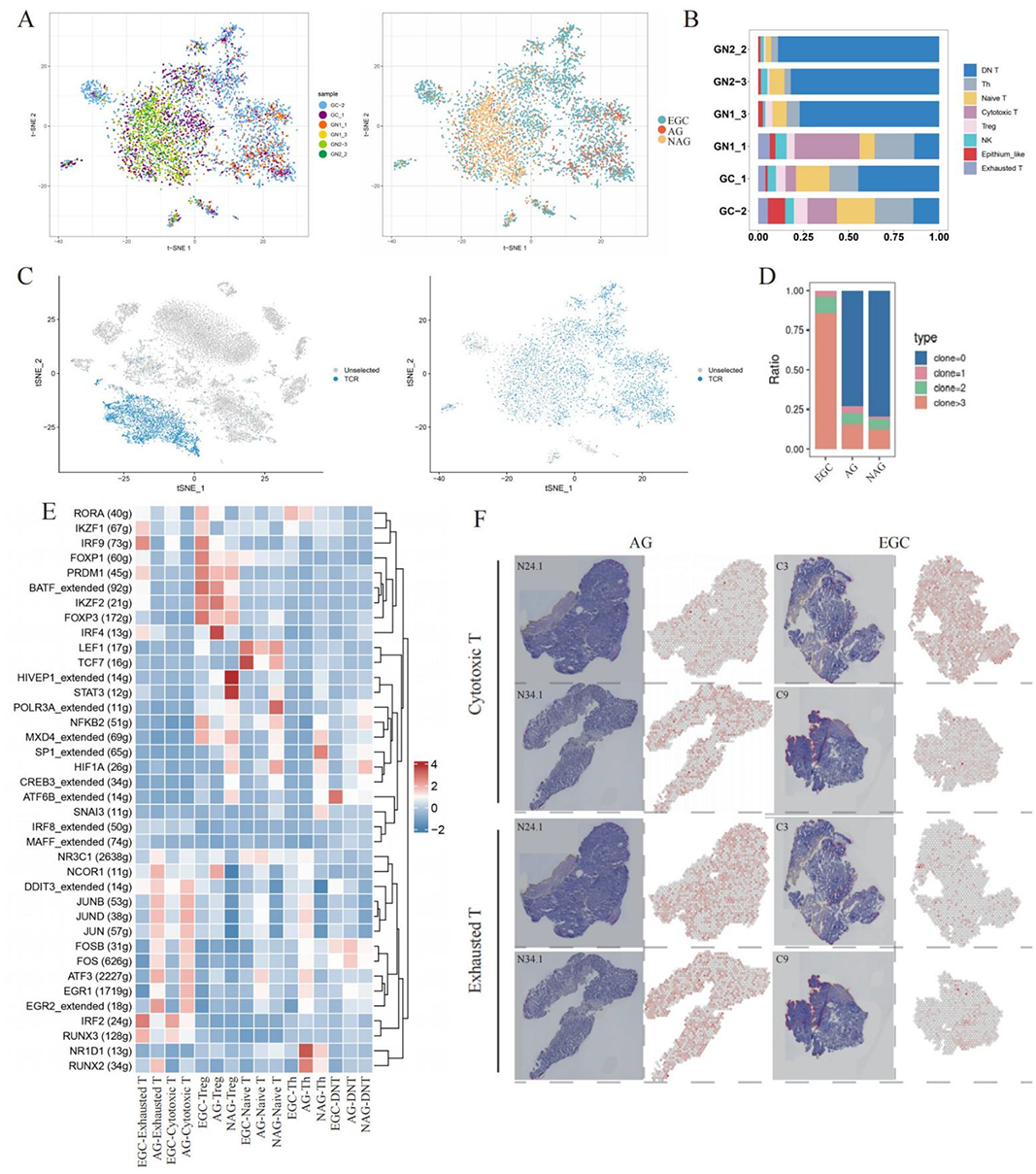
The features of T cells, related to Figure 3. (A) The pathological information of patients used in this study. (B) The TSNE feature plots of marker genes for identified celltypes. (C) The cell proportions of identified celltypes among 6 patients used for scRNA-seq. (D) The expression of TFF2 among NAG, AG and EGC groups. (E) Average proportion of each spot in identified celltypes. (F) The celltype location in spatial images (C8, N21-1 and N22-1) of fibroblasts, parietal cells, endothelium, mast cells, endocrine cells and pericytes.

**Supplementary Figure 8.**
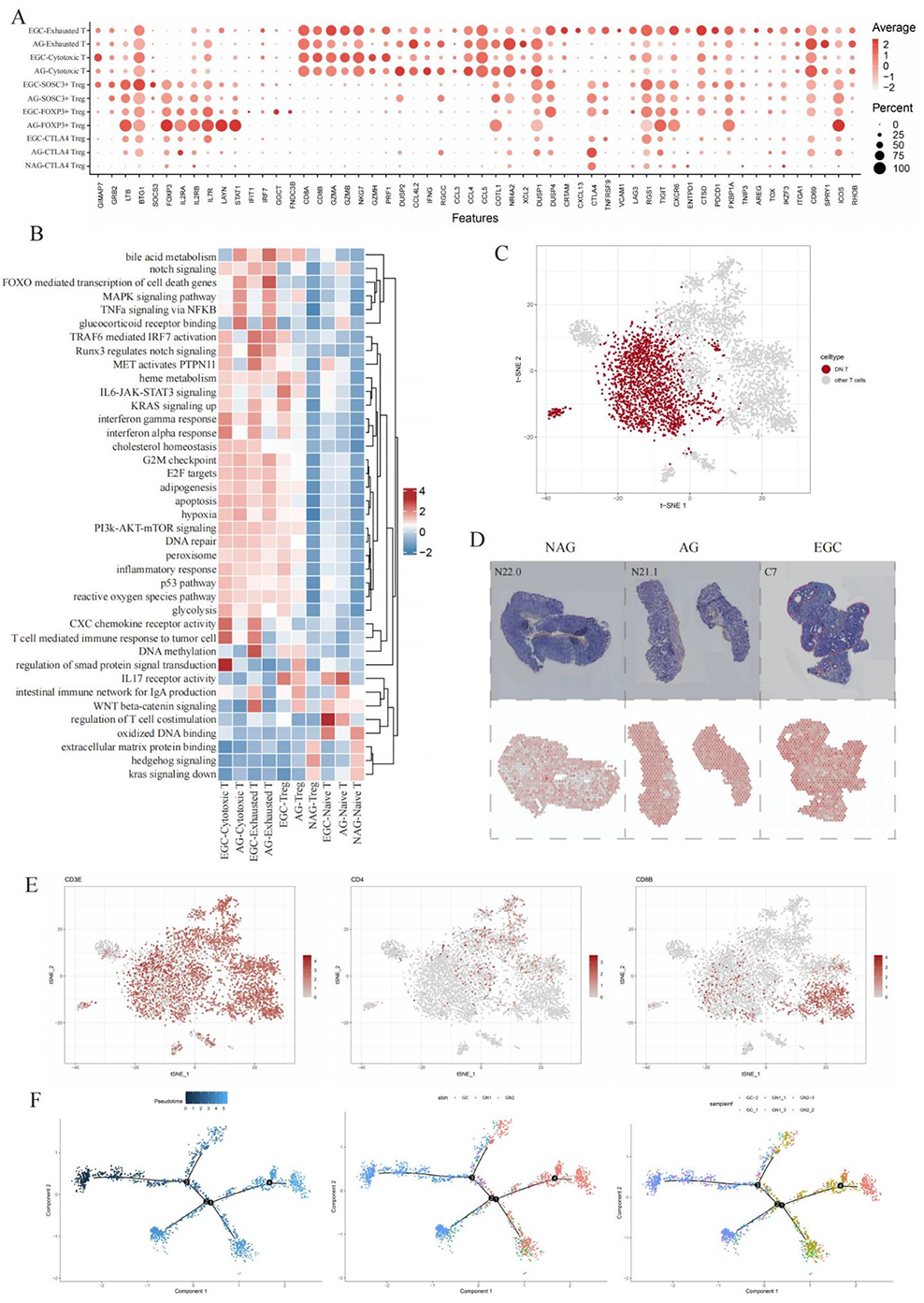
The features of specific subclusters of T cell and identification of DNT cells, related to Figure 3. (A) The expression of selected genes showed in different clusters of Tregs, cytotoxic and exhausted T cells grouped by NAG, AG and EGC. (B) The pathways enriched in different clusters of Tregs, cytotoxic and exhausted T cells grouped by NAG, AG and EGC. (C) The highlighted DNT cells among all T cells. (D) Spatial location of DNT cells in N22.0, N21.1 and C7. (E) The definition of DNT cells with rare expression of CD4 or CD8 markers. (F) The different states used by time trajectory analysis for NDT cells.

**Supplementary Figure 9.**
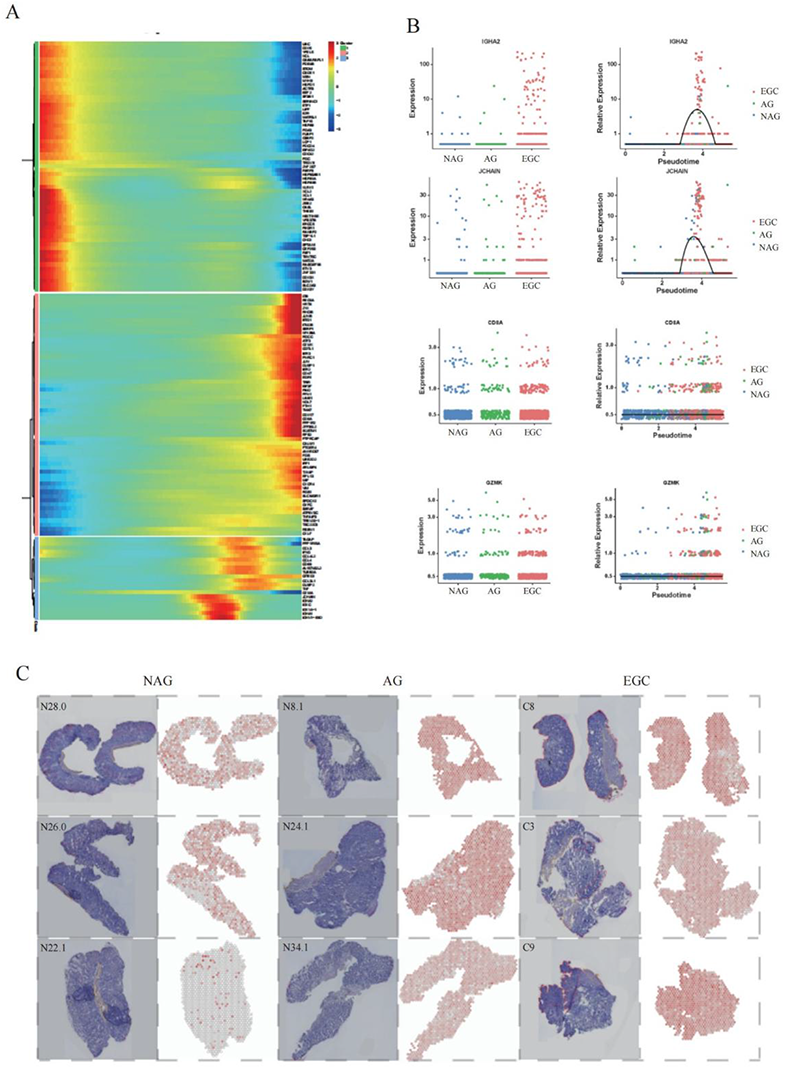
The DEGs of DNT cells, related to Figure 3. (A) The DEGs followed the changes of time trajectory. (B) The gene expression along with the changes of time trajectory. (C) Spatial location of DNT cells in NAG (N28.0, N26.0 and N22.1), AG (N8.1, N24.1 and N34.1) and EGC (C8, C3 and C9) groups.

**Supplementary Figure 10.**
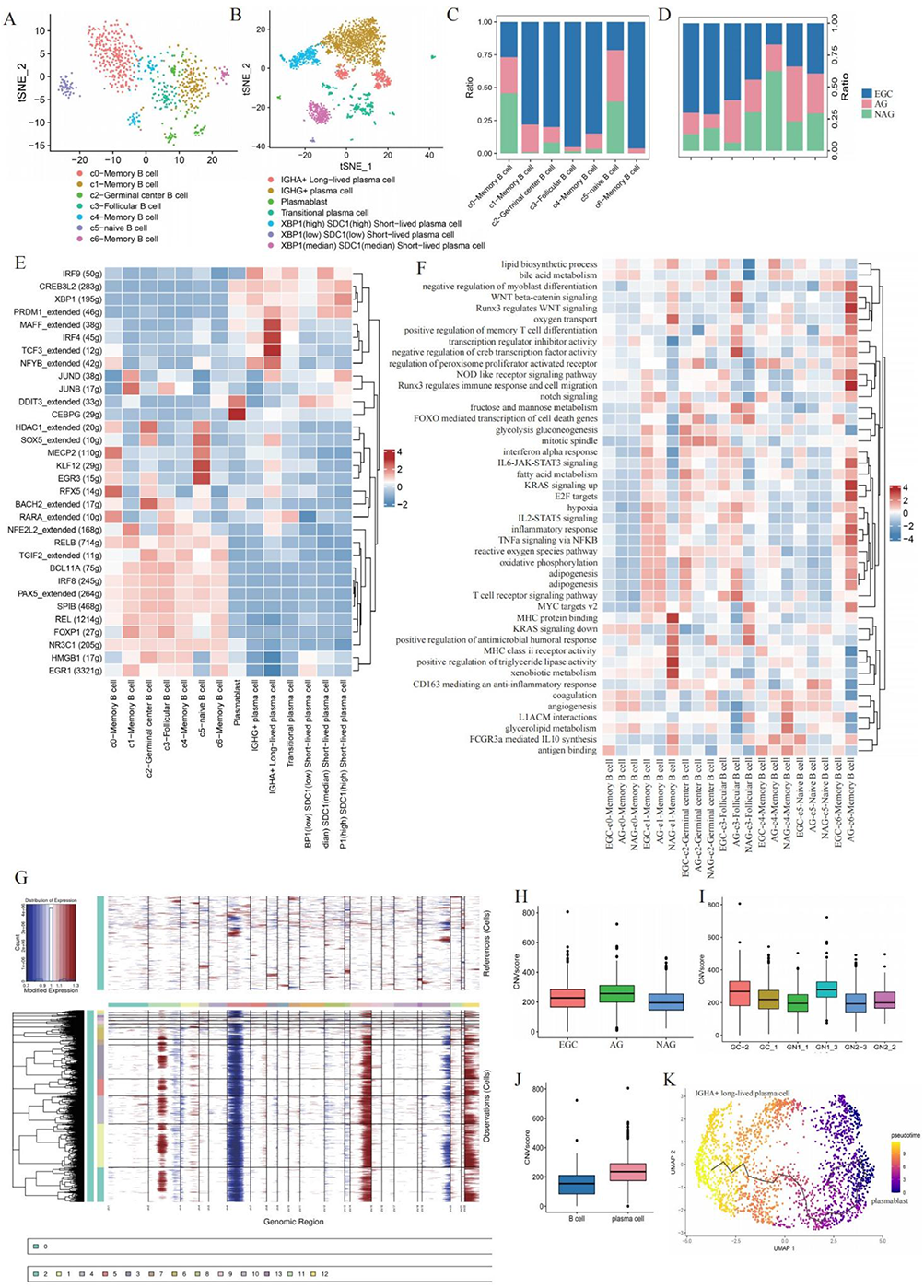
The pathway and transcriptional activities of subclusters in B cells, related to Figure 4. (A) The TSNE plot of subclusters in B cells. (B) The TSNE plot of subclusters in plasmas. (C) The proportion of subclusters in B cells. (D) The proportion of subclusters in plasmas. (E) The transcriptional factors in subclusters of B cells and plasmas. (F) The pathway activities in subclusters of B cells and plasmas. (G) The CNV of B cells estimated by inferCNV. (H) The CNV scores among groups, (I) samples and between (J) B cells and plasmas. (K) The time trajectory of subclusters in plasmas.

**Supplementary Figure 11.**
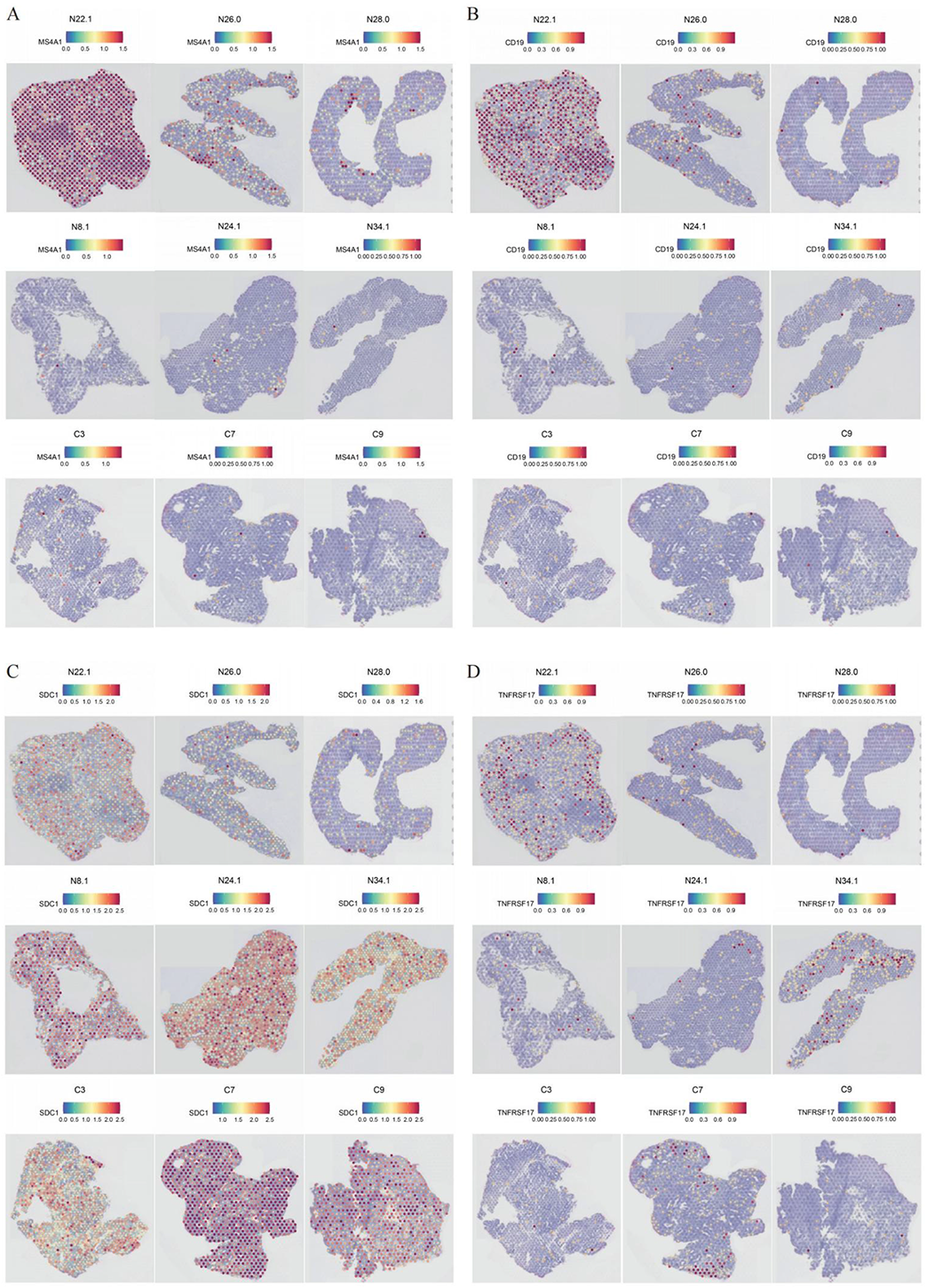
Spatial location and expression for marker genes of B cells, related to Figure 4. (A) The spatial location and expression for MS4A1 (CD20), (B) CD19, (C) SDC1, and (D) TNFRSF17 among all 12 samples.

**Supplementary Figure 12.**
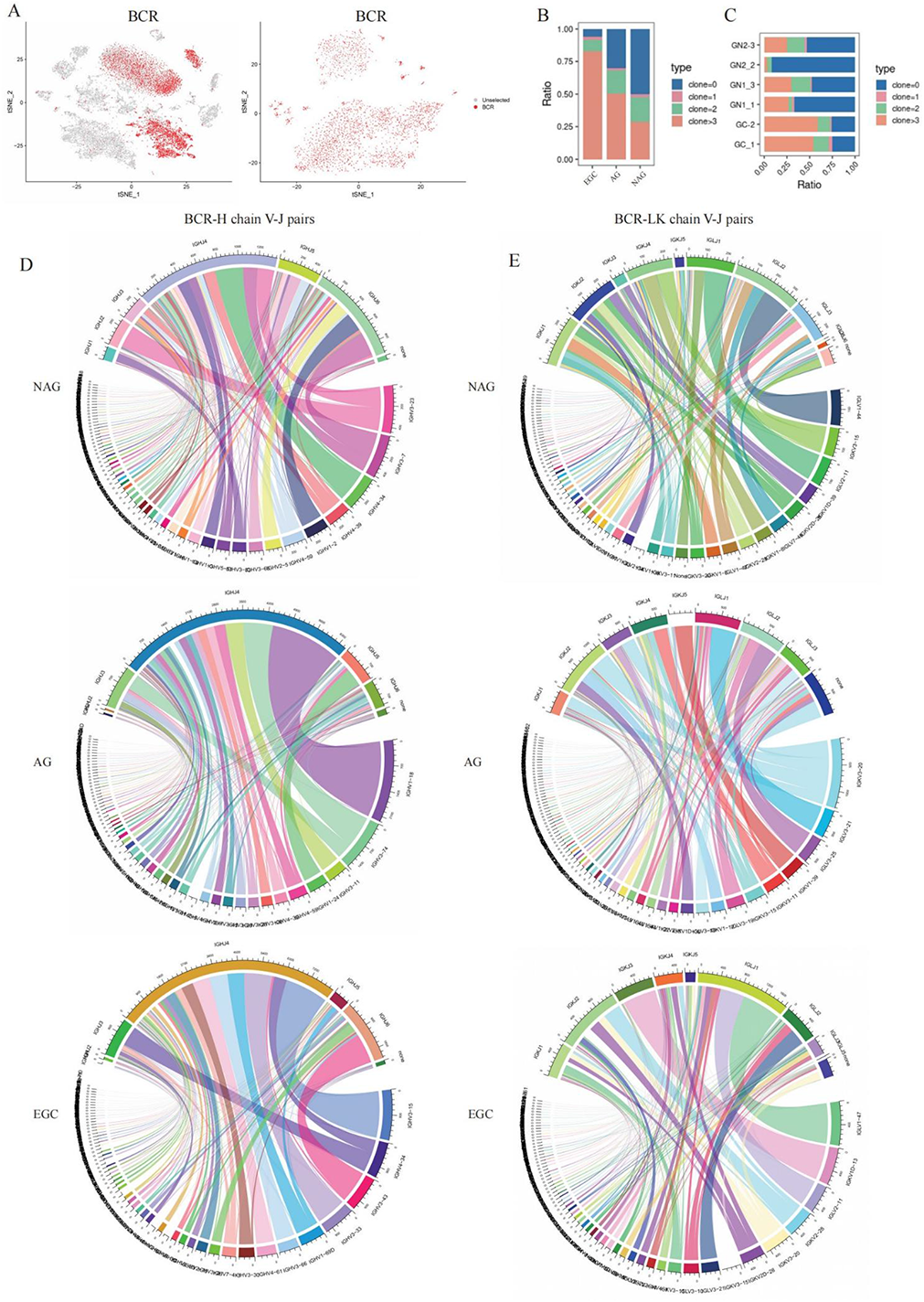
The BCR changes of B cells with the progression of EGC, related to Figure 4. (A) The cells with BCR were highlighted on all cells and B cells. (B) The BCR diversity among NAG, AG and EGC groups. (C) The BCR diversity among 6 samples. (D) The V-J pairs of BCR heavy chains (H represented the heavy chains) and (E) light chains (LK represented the two light chains)

**Supplementary Figure 13.**
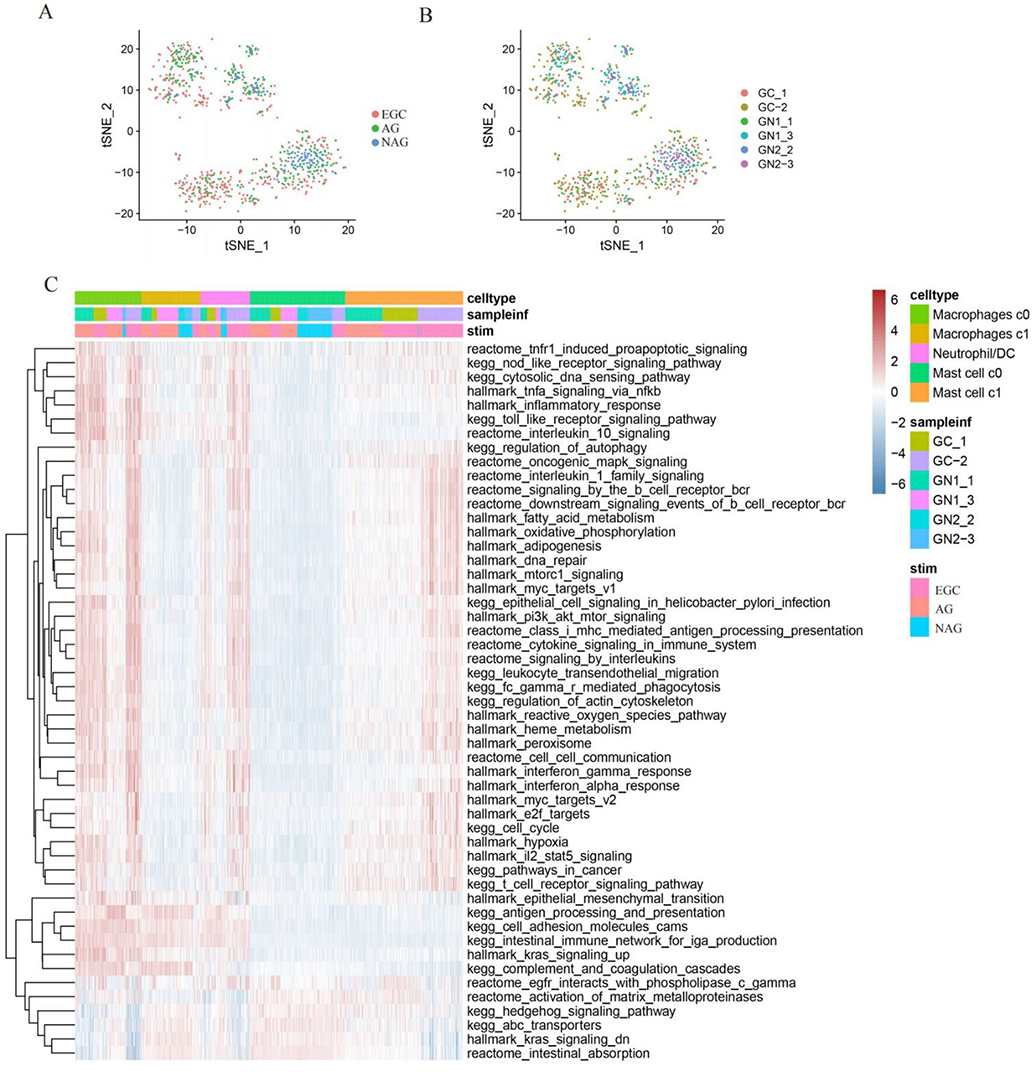
The pathway activities of myeloid cells, related to Figure 5. (A) The TSNE plot of myeloid cells among 3 groups. (B) The TSNE plot of myeloid cells among 6 samples. (C) The pathway activities of different subclusters of myeloid cells.

**Supplementary Figure 14.**
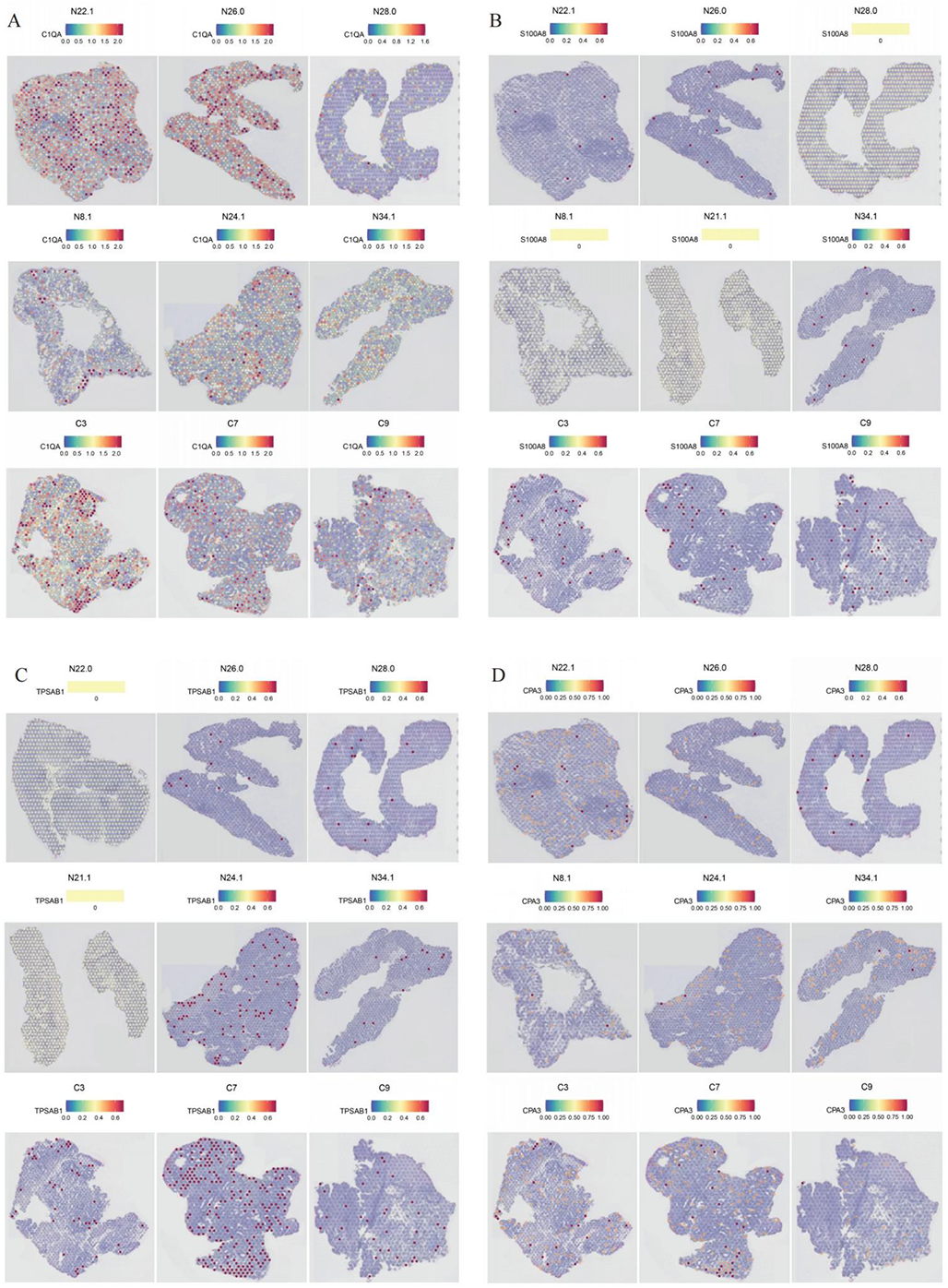
Spatial location and expression for marker genes of myeloid cells, related to Figure 5. (A) The spatial location and expression for C1QA, (B) S100A8, (C) TPSAB1, and (D) CPA3 among all 12 samples.

**Supplementary Figure 15.**
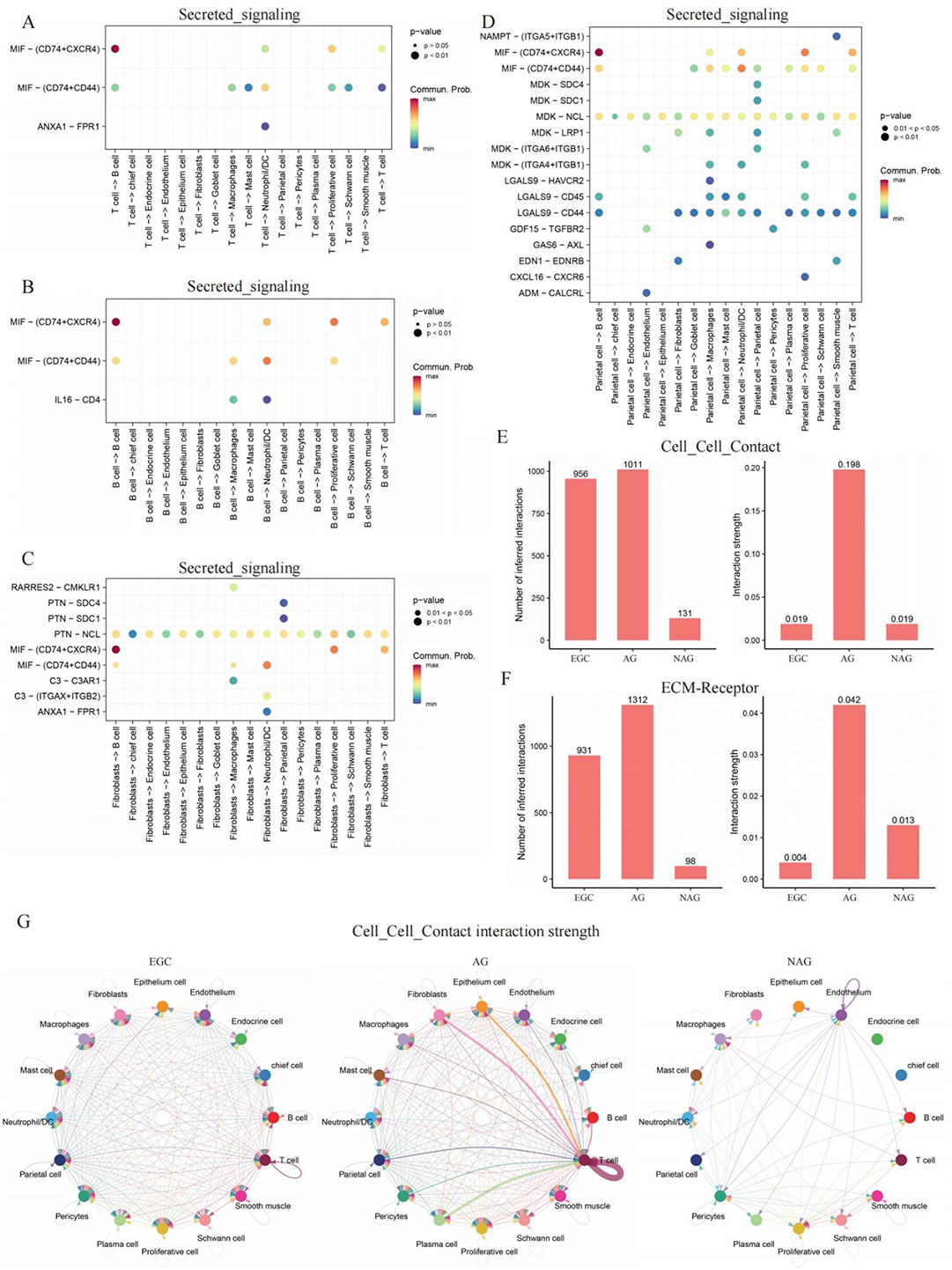
The cellular interaction of secreated signaling and cell-cell contact signaling among specific cells, related to Figure 6. (A) The cellular interaction between T cells and other cells about secreated signaling. (B) The cellular interaction between B cells and other cells about secreated signaling. (C) The cellular interaction between fibroblasts and other cells about secreated signaling. (D) The cellular interaction between parietal cells and other cells about secreated signaling. (E) The interaction numbers and strengths of cell-cell contact signaling and (F) ECM signaling. (G) The cellular interaction strengths of cell-cell contact signaling for each pair among NAG, AG and EGC groups.

**Supplementary Figure 16.**
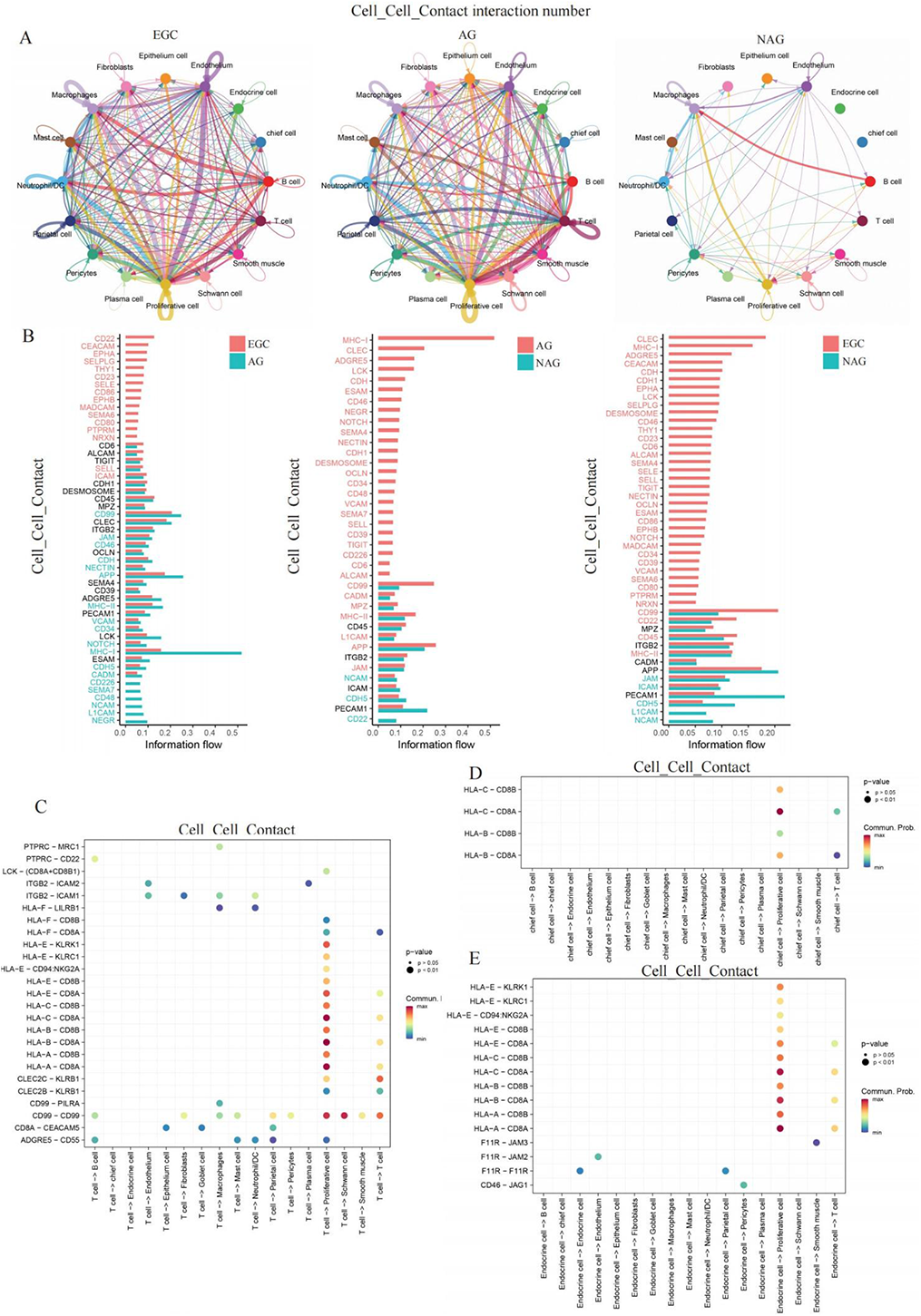
The cellular interaction of cell-cell contact signaling among specific cells, related to Figure 6. (A) The cellular interaction numbers of cell-cell contact signaling for each pair among NAG, AG and EGC groups. (B) The interaction involved pathways between AG and EGC (left), NAG and EGC (median), as well as NAG and AG (right) of cell-cell contact signaling. (C) The cellular interaction between T cells and other cells about cell-cell contact signaling. (D) The cellular interaction between chief cells and other cells about cell-cell contact signaling. (E) The cellular interaction between endocrine cells and other cells about cell-cell contact signaling.

**Supplementary Figure 17.**
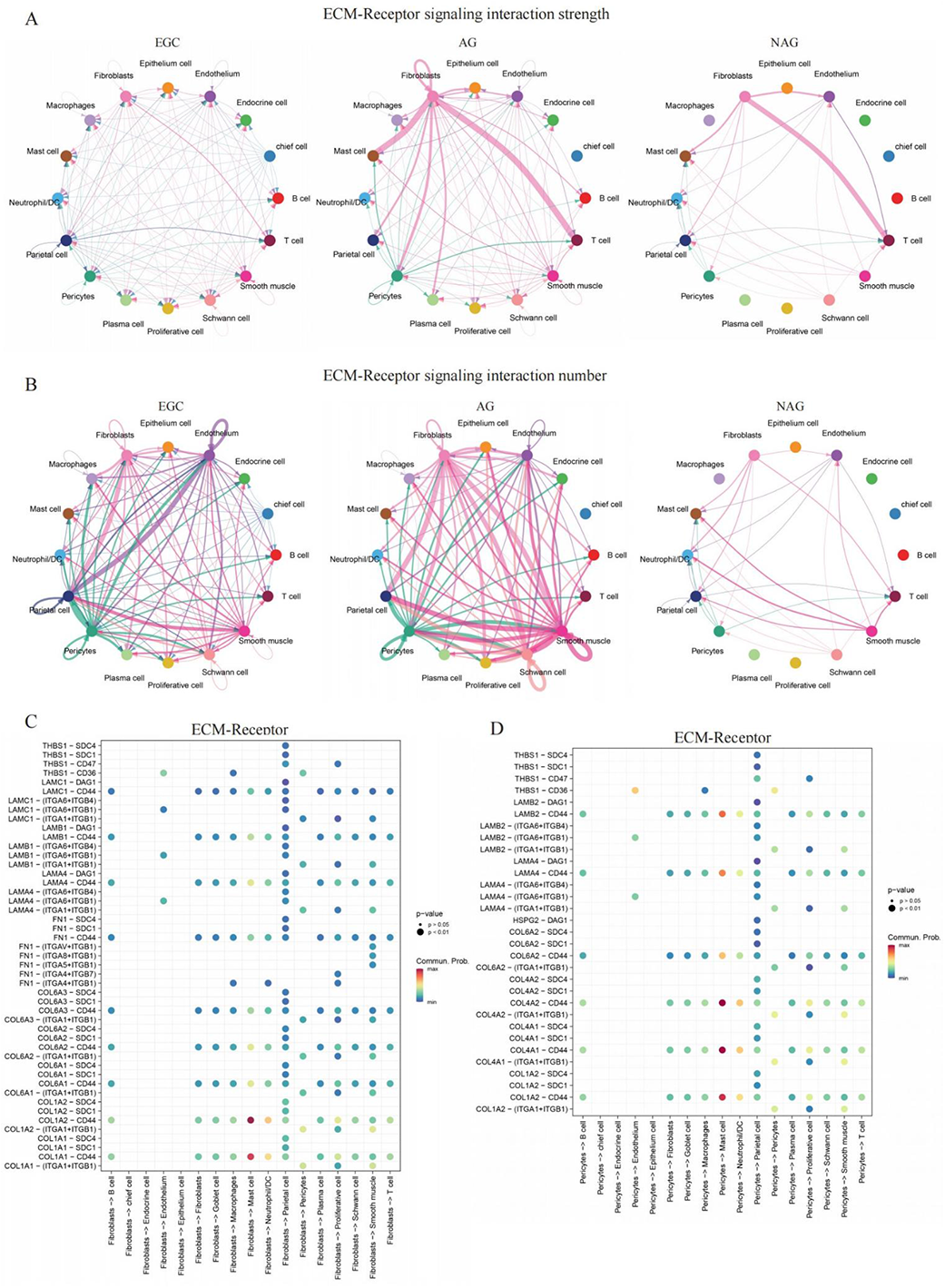
The cellular interaction of ECM signaling among specific cells, related to Figure 6. (A) The cellular interaction strengths of ECM signaling for each pair among NAG, AG and EGC groups. (B) The cellular interaction numbers of ECM signaling for each pair among NAG, AG and EGC groups. (C) The cellular interaction between fibroblasts and other cells about cell-cell contact signaling. (D) The cellular interaction between pericytes and other cells about cell-cell contact signaling.

